# Ultrahigh-throughput discovery of modified aptamers as specific and potent enzyme inhibitors

**DOI:** 10.1101/2024.08.16.608213

**Authors:** Claire Husser, Janis Hoetzel, Roger Cubi, Isabelle Lebars, Leon Kraus, Carmelo Di Primo, Stephanie Baudrey, Ewgenij Proschak, Bruno Kieffer, Beatrix Suess, Michael Ryckelynck

## Abstract

Enzymes are instrumental to life and key actors of pathologies, making them relevant drug targets. Most enzyme inhibitors consist of small molecules. Although efficient, their development is long, costly and can come with unwanted off-targeting. Substantial gain in specificity and discovery efficiency is possible using biologicals. Best exemplified by antibodies, these drugs derived from living systems display high specificity and their development is eased by harnessing natural evolution. Aptamers are nucleic acids sharing functional similarities with antibodies while being deprived of many of their limitations. Yet, the success rate of inhibitory aptamer discovery remained hampered by the lack of an efficient discovery pipeline. In this work, we addressed this issue by introducing an ultrahigh-throughput strategy combining *in vitro* selection, microfluidic screening and bioinformatics. We demonstrate its efficiency by discovering a modified aptamer that specifically and strongly inhibits SPM-1, a beta-lactamase that remained recalcitrant to the development of potent inhibitors.

**Graphical Abstract:** 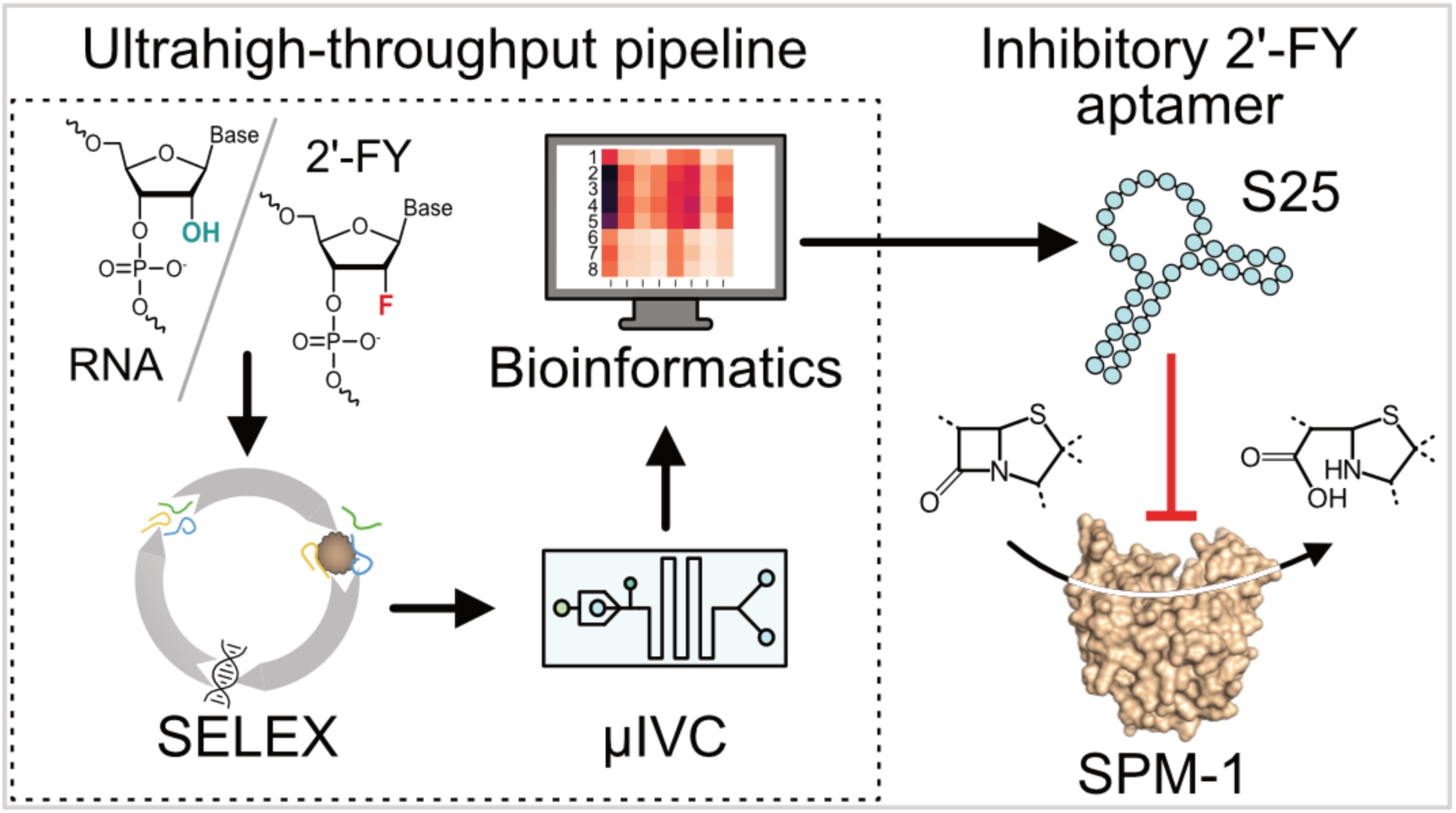

## Introduction

Whether confined within the cell or secreted, enzymes are instrumental actors of life through their pleiotropic involvement in processes as diverse as cell growth and division, metabolism, gene expression and control, motility, or even organism morphogenesis just to name a few examples^1^. Besides their key functions in cell and organism homeostasis, enzymes are also major players in the appearance and progression of diseases (e.g., genetic disorders and cancer) and infectious processes. Therefore, being able to specifically inhibit an enzyme not only offers an exquisite way of studying its function, but this also paves the way for the development of efficient therapeutics.

Most enzyme inhibitors consist of small chemicals of natural (especially toxins^2^ and antibiotics^3^) or synthetic origin. Indeed, a large palette of substrate analogues^4^, suicide substrates^5^ and non-competitive inhibitors^1^ have now been discovered to (ir)reversibly inhibit target enzymes. Although chemical modeling and chemoinformatics can assist drug development^6^, discovering specific and potent chemical inhibitors is a long and tedious process requiring important synthesis, screening and testing efforts. Moreover, small molecules can lack specificity and have side effects. Biologicals represent an attractive alternative to reduce the labor and the development cost of new inhibitors while improving both discovery rate and drug specificity^7^. This new paradigm relies on the use of cells or molecules (e.g., hormones, peptides, antibodies, proteins and nucleic acids) originating from a biological system. A key advantage of biologicals lies in their programmability and evolvability. Indeed, living systems have the capacity to encode biological molecules and store instruction modes within a nucleic acid (usually deoxyribonucleic acid, DNA). Mutating this polymer and selecting organisms or biological polymers with improved fitness form the basis of the evolution by natural selection as originally theorized by Charles Darwin. Whereas this process usually operates over extended periods of time in nature, some immune cells acquired the ability to rapidly accumulate mutations in antibody-coding genes. This hypermutation activity drives the so-called affinity maturation process allowing the rapid selection of cells producing best-performing antibodies against nearly any target^8^. This concept is now massively exploited for the discovery of antibodies with a wide application spectrum, making them the largest class of biologicals^9,10^, especially as enzyme inhibitors^11^. Although several therapeutic antibodies are now on the market^9^, their development is long, it requires the use of animals, long screening steps are needed to identify the cells producing inhibitory antibodies, their production is costly and can be prone to batch variability.

Nucleic acids are another promising class of therapeutics currently best represented by RNA vaccines, antisense oligonucleotides and siRNA mimetics^12,13^. These biologicals are mainly developed *in vitro* (avoiding the use of animals and allowing the use of toxic compounds), they have a low immunogenicity, they can be produced in large scale using enzymatic or solid phase synthesis and they can easily be chemically modified. This class of biologicals is completed by aptamers, also referred to as “chemical antibodies” because of functional similarities^14^. Indeed, these oligonucleotides do not simply recognize target molecules through base-pairing, but they instead adopt a complex three-dimensional structure enabling them to specifically recognize their target by shape complementary, the establishment of hydrogen bonds, electrostatic or van der Waals interactions. A great advantage of RNA aptamers is the possibility of isolating them *de novo* by *in vitro* selection using methodologies collectively known as systematic evolution of ligands by exponential enrichment, SELEX in short^15,16^. Briefly, a pool of 10^15^ different random sequences is challenged to interact with a target prior to partitioning them away from the bulk through extensive washes. Performing iterative rounds then enables to progressively enrich the pool in highly specific binders. A plethora of aptamers has now been discovered and two of them have been approved by the FDA as treatments for age-related macular degeneration (Macugen, a VEGF inhibiting aptamer^17^ and Izervay, a complement C5 inhibiting aptamer^18^). Both aptamers bind their targets at sites inhibiting downstream recognition by a molecular partner (receptor or enzyme). Transposed to the search of enzyme inhibitors, *in vitro* selection offers the great advantage of its blindness, leaving the door open to the identification of any type of inhibitor (competitive, non-competitive, uncompetitive). Several enzyme inhibitory aptamers have also been described^19–21^, although real world applications are still lacking behind their potential. A current important limitation stems from the way aptamers are searched. Indeed, in SELEX candidates are primarily selected for their binding capacity, a feature not directly correlated with an inhibit capacity. Consequently, the identification of an inhibitory aptamer within a SELEX-enriched pool requires tedious post-selection processing often restricted to a few dozen molecules, limiting the success rate of campaigns aiming at identifying such molecules. This parallels our previous observations establishing that SELEX-derived aptamers are often suboptimal for functions more advanced than binding (e.g., fluorescence emission^22,23^ and gene expression control^24,25^). Such a limitation can nevertheless be overcome by high-throughput screening methodologies like the microfluidic-assisted *in vitro* compartmentalization used in tandem with high-throughput sequencing (µIVC-seq) that we originally introduced for the efficient discovery light-up aptamers^26–28^.

In the present work, we combined the use of SELEX in tandem with µIVC-seq and deep bioinformatic analyses to set-up an ultrahigh-throughput pipeline enabling the efficient identification of potent enzyme inhibitors suited for applications in challenging biological environments. To test this novel approach, the São Paulo metallo-beta-lactamase 1 (SPM-1) was chosen as the first target. SPM-1 is naturally expressed by *Pseudomonas aeruginosa* and contributes to multidrug-resistance of the bacteria by antibiotics hydrolysis^29,30^. As part of the dawning antibiotic resistance crisis^31^, SPM-1 is a relevant target for drug development. While some drugs are known to inhibit serine beta-lactamases, no specific inhibitor for metallo-beta-lactamases has been discovered yet^32^. We therefore chose this enzyme as a target to evaluate the capacity of our pipeline to discover a specific and potent inhibitor. Importantly, future applications in challenging biological environments place the nuclease resistance of such an aptamer at the forefront of expected features. This issue can be addressed by exploiting chemical modifications known to endow aptamers with extended stability and/or capacity to establish additional interactions with target molecules^12,14,33^. However, to limit possible difficulties, the modification should better be included during aptamer selection rather than being introduced in a second development step. We therefore adapted our pipeline to the screening of libraries expressed as RNA or as oligonucleotides made of 2’-fluorinated modified pyrimidines (2’-FY), a chemistry known to strongly increase the stability of nucleic acids^34^. Using this new strategy compatible with any enzyme, we efficiently identified an aptamer endowed with an impressive half-life in serum, high affinity, specificity and inhibition capacity of the recalcitrant SPM-1 metallo-beta-lactamase. Moreover, using bioinformatics we were able to monitor for the first time how a pool adapts to the introduction of a new chemistry, while biophysical characterization demonstrated that, beyond their expected impacts, modifications can have additional and hard to predict effects supporting the need of introducing them early in the development pipeline.

## Results

### Isolation of SPM-1 inhibitory RNA aptamers

To isolate an SPM-1 inhibitory aptamer, a pool of 1.2 x 10^15^ molecules with a region of 40 randomized nucleotides (Supplementary Fig. 1) was subjected to an *in vitro* selection (SELEX) to pre-enrich the pool in SPM-1 binders prior to isolating the most potent inhibitors by microfluidic-assisted ultrahigh-throughput functional screening (Fig. 1a and b). We started by expressing the pool in RNA chemistry and challenged it to interact with an immobilized His-tagged form of SPM-1 (Fig. 1a, left). An enrichment of target binders was already distinguishable starting from round R5 of SELEX and it kept on increasing until round R7 (Fig. 1c). The relative fraction of SPM-1-bound RNA increased from 0.18% in the first round to nearly 10% in round R7. The amount of RNA binding to target-free beads did not exceed 0.4%, suggesting a specific binding to the target protein. Moreover, inhibition assay revealed that, whereas the starting pool did not yield any measurable inhibition of SPM-1 activity (Fig. 1d), the pool enriched after round R7 displayed a slight but reliable effect on the enzyme indicating that a subset of this pool not only bound to SPM-1, but also interfered with its activity.

**Fig. 1.**
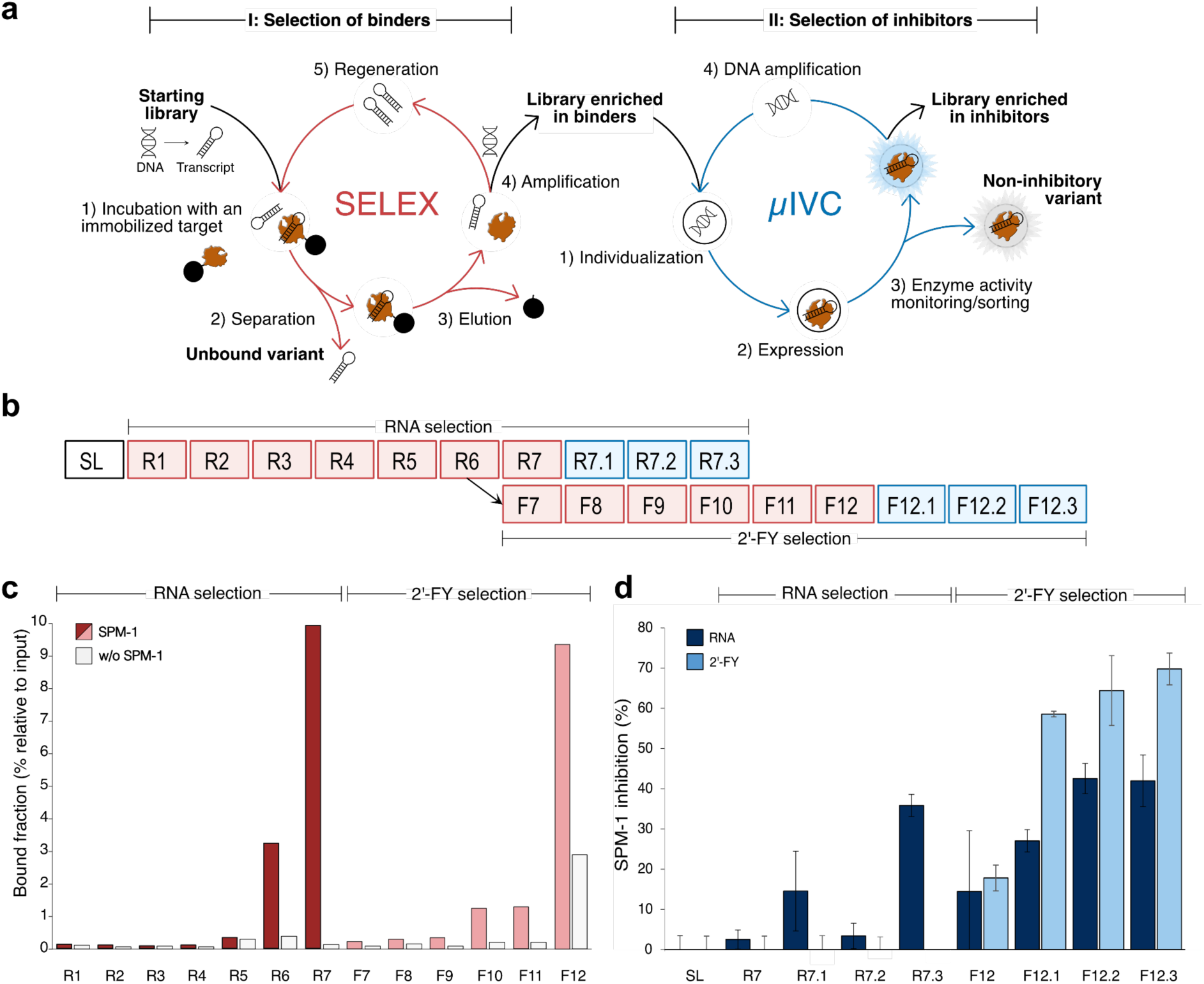
Dual selection for binding and inhibition capacity. **a**, Schematic of the dual selection approach. The RNA library was first enriched in binders using SELEX. Briefly the library was incubated with the target protein immobilized on beads prior to washing away unbound oligonucleotides and recovering only those target-bound ones. Variants of interest were then eluted, and the library was regenerated prior to subjecting it to another round of selection. After several rounds, the binder-enriched library was screened for activity using ultrahigh-throughput microfluidic screening (µIVC). Genes of the enriched library were first individualized and amplified in droplets containing a PCR mixture, prior to being fused to an *in vitro* transcription mixture supplemented with the target enzyme. The amplified DNA was transcribed in the presence of SPM-1 during a second incubation step. Droplets were then reinjected into an integrated chip where they received a controlled amount of SPM-1 substrate, they were incubated for a defined time at controlled temperature and sorted according to their in beta-lactamase activity. **b**, Experimental selection workflow. The starting library (SL) was expressed as RNA and subjected to 7 rounds of SELEX (red boxes, R1-R7) followed by 3 rounds of µIVC (blue box). The selection process was next pursued in 2’-fluorinated chemistry (2’-FY) starting from round R6. 6 rounds of SELEX (F7-F12) were performed while expressing libraries in 2’-FY, followed by 3 rounds of µIVC. **c.** Elution profile of the oligonucleotide bound fraction during SELEX experiments. The fraction of oligonucleotide eluted from SPM-1-coupled beads (red bars) or empty beads (negative control, white bars) is given as the percentage of the total input RNA used in each round. **d**, Enrichment profile of the libraries in SPM-1 inhibitory aptamers. Libraries recovered after each round of screening (R7.1-R7.3 and F7.1-F7.3) were either expressed in RNA (dark blue) or in 2’-FY (light blue) and their capacity to inhibit SPM-1 activity was assessed using CCF2-FA assay. Values are the mean of 3 independent experiments and the error bars correspond to the standard deviation. SPM-1 inhibition was normalized by the condition without inhibitors. Error bars were cut upon axis interception for an illustrative purpose.

To isolate the most potent SPM-1 inhibitors, we tailored our µIVC-seq functional screening pipeline to individually evaluate the inhibition capacity of each variant and conserve only the most efficient ones (Fig. 1a, right). Briefly, the DNA sequences coding for the variants enriched after the round R7 of SELEX were individualized into water-in-oil droplets where they were PCR-amplified prior to being *in vitro* transcribed in the presence of SPM-1. Upon incubation, a controlled amount of CCF2-FA^35^, a fluorescent substrate of beta-lactamase (Supplementary Fig. 2a), was delivered to each droplet by picoinjection. To maximize the chance of isolating aptamers of interest, their inhibition capacity was assayed after a short incubation time (*i.e.*, 5 minutes) at biologically relevant temperature (*i.e.*, 37°C) and with a substrate concentration (*i.e.*, 20 µM) expected to saturate enzyme (SPM-1 has a *K_M_* of ∼2.5 µM for CCF2-FA^36^). To do so, we used an integrated device in which a picoinjector was directly connected via a delay line to an analysis and sorting module and we held the device at 37°C throughout the process (Supplementary Fig. 2b). Upon only 3 rounds of µIVC (rounds R7.1-R7.3, Supplementary Fig. 3a), the pool was strongly enriched in potent inhibitors (36% inhibition of SPM-1 activity, Fig. 1d).

Upon libraries sequencing, we identified 40 clades of highly similar motifs appearing, even transiently, at a > 0.5 % frequency throughout the whole process (Supplementary Fig. 4a). Some clades (*e.g.*, clades 9 and 22) were found to occur at constant but rather low frequencies (Supplementary Fig. 4b). On contrary, other clades rapidly depleted after 5 to 6 rounds of SELEX (*e.g.*, respectively clades 34-38 and 21) prior to being replaced by several new clades (*e.g.*, clades 17, 25, 28, 32, 42 and 47) starting from round R5 (Fig. 2a and Supplementary Fig. 4b). Finally, starting from round R7 clades 7, 8 and 16 started to be enriched and maintained until the end of the process (*i.e.*, round R7.3). Interestingly, clade distribution remained constant throughout µIVC screening steps (Fig. 2a), while sequence diversity dropped from > 130,000 unique sequences in Round-7 to merely 4,400 unique sequences in round R7.3 (Supplementary Fig. 4c). This sequence enrichment nicely parallels the increasing inhibition capacity of the corresponding libraries (Fig. 1d). All together, these observations highlight that µIVC screening predominantly worked by depleting libraries of the least efficient sequences without major remodeling of clade content.

**Fig. 2.**
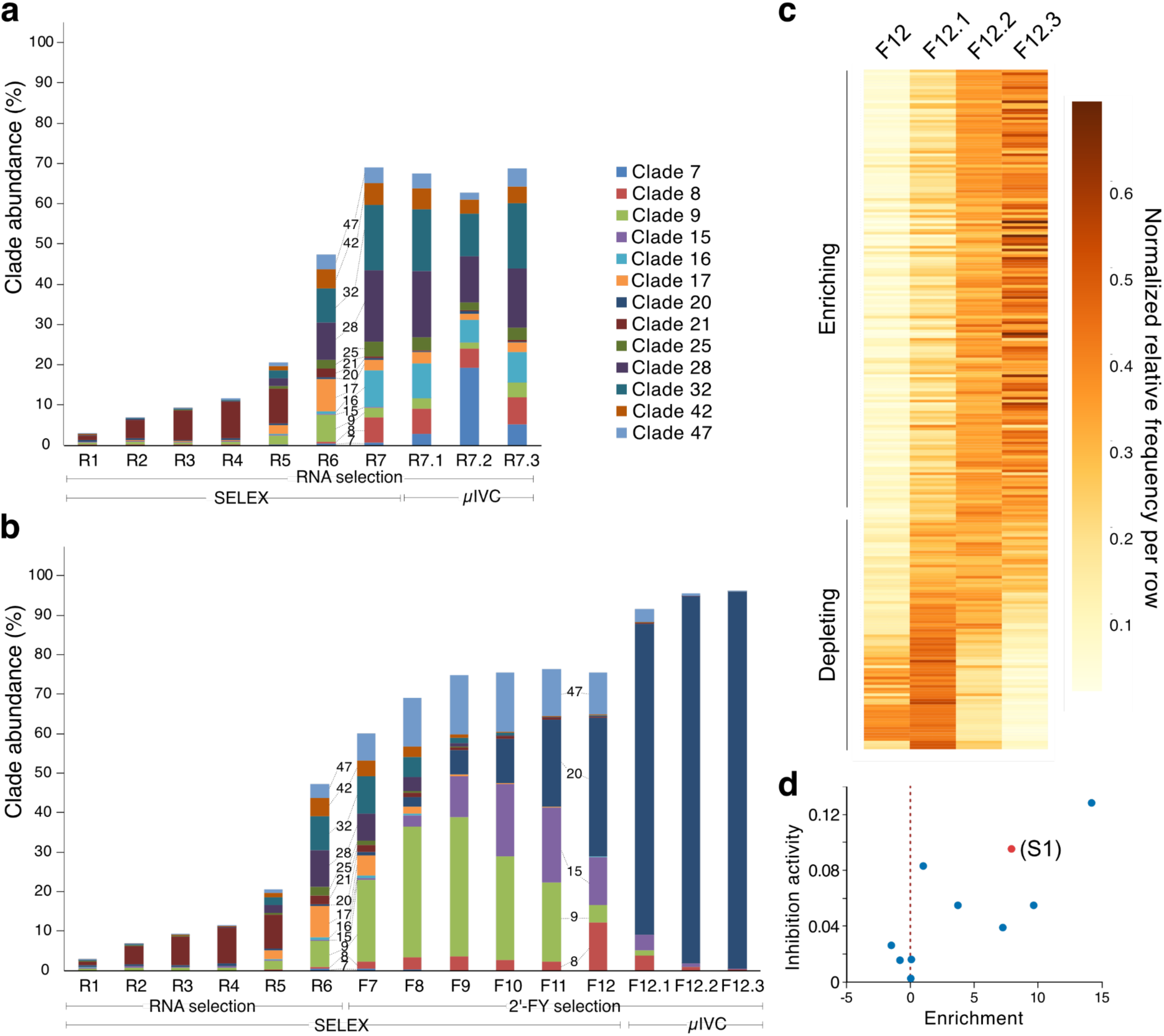
Monitoring of sequence accumulation and depletion throughout the selection process. **a**, Evolution of clade identity and abundance during RNA-based selections. The relative abundances of the most representative clades is indicated for each round of SELEX (R1-R7) and µIVC screening (R7.1-R7.3). **b**, Evolution of clade identity and abundance during RNA-based (R series) and later by 2’-FY-based (F series) selections. The relative abundances of the most representative clades is indicated for each round of SELEX during which libraries were expressed as RNA (R1-R6) and then in 2’-FY (F7-F12) followed by µIVC screening (F12.1-F12.3). Each bar represents a selection round, a different color was attributed to each clade and their relative abundance is also encoded by the height of the bar. **c**, Enrichment of the sequences contained in clade 20. The frequency of each sequence displaying a cumulative frequency higher than 0.1 was computed for each round and data were normalized by row. Sequences were finally ordered by their enrichment allowing enriching sequences to be discriminated from the depleting ones throughout the process. **d**, Evaluation of the inhibition capacity of a selection of variants. 10 representative aptamers of clade 20 with different enrichment trends were tested for their capacity to inhibit SPM-1 activity.

### Searching for nuclease resistant SPM-1 inhibitory aptamers

As future applications would require the aptamer to remain stable in challenging extracellular environments, we evaluated to what extent aptamers contained in the enriched libraries would preserve their function when transcribed as 2’-fluorodeoxypyrimidines (2’-FY), a chemistry known to substantially protect aptamers from rapid degradation^34^. Yet, expressing libraries R7 to R7.3 as 2’-FY molecules did not yield any detectable inhibition of SPM-1 activity (Fig. 1d). Therefore, the R6 pool (expected to still contain a significant sequence diversity) was expressed as 2’-FY aptamers and subjected to additional rounds of SELEX and µIVC screening (Fig. 1b). As expected, only a small subfraction of aptamers preserved a capacity to interact with SPM-1 as protein-bound fraction dropped down to 0.23%, but 6 rounds (F7 to F12) of SELEX were sufficient to restore this fraction to 9.3% (Fig. 1c). As before, the last SELEX-enriched library was further subjected to 3 rounds of µIVC screening (Supplementary Fig. 3b) and yielded F12.3 library endowed with much higher inhibition capacity of SPM-1 activity (to nearly 70%, Fig. 1d). Interestingly, although still functional when expressed as RNA, F12.3 library displayed optimal performances upon transcription using 2’-FY nucleotides. This better adaptation to 2’-FY chemistry was also apparent at the sequence level since only 4% of them were found common between the pools obtained from round R7 and F12 (Supplementary Fig. 4f). Interestingly, clade 9 first benefited from the use of 2’-FY chemistry and tended to be the dominant species after only 3 rounds of SELEX (Fig. 2b and Supplementary Fig. 4e), prior to being progressively supplanted by clades 8, 15 and 20 that became the main species found after the last round of SELEX (F12). As no change in selection pressure was introduced during the SELEX, this change in library composition was attributed to competition between aptamers. µIVC functional screening led again to the depletion of nearly 98% (going from ∼117,000 sequences in F12 to ∼ 2,700 in F12.3, Supplementary Fig. 4d) of the sequences, leaving only clade 20 (> 95 % of the sequences) as the dominant species at the end of the process (Fig. 2b and Supplementary Table1). Noteworthy, µIVC screening also enabled the efficient removal of shorter-sized molecules (Supplementary Fig. 5) known to frequently accumulate and poison SELEX experiments^37^.

Further evaluation of clade 20 identified 229 different sequences displaying significant appearance frequency, of which around two thirds displayed continuous enrichment throughout the process while the remaining third tended to deplete (Fig. 2c). Interestingly, the functional evaluation of a few variants also indicated that the inhibition capacity of an aptamer somehow correlates with its tendency to get enriched (Fig. 2d). Deeper sequence analysis identified S1 as the consensus sequence of clade 20 (Supplementary Table 2) as well as the most abundant (*i.e.*, 15%) variant in library F12.3 (Supplementary Fig. 6), and an efficient inhibitor of SPM-1 (Fig. 2d).

### Optimization of SPM-1 inhibitory aptamer

Analyzing S1 with Vienna Package RNAfold generated a simple secondary structure model (Fig. 3a) in which a long P1 stem forms between residues 8-20 (encompassing the 5’ constant region) and 42-55, connected to a shorter P2-L2 stem-loop (residues 31-41) *via* a large internal loop (J1-2, residues 21-30). Careful analysis of sequencing data of the three screening steps enabled to map regions tolerating or not variations by computing a functional prediction score for each position and allowed to delineate the functional part of the aptamer to its apical region (Fig. 3a). Indeed, some variations were generally accepted in P1, consistent with the possibility of deleting the A47 bulging residue and replacing most of the P1 helix by an alternative stem of different sequence (mutant S1.5) with minimal impact on aptamer functionality (Fig. 3b). On the contrary, the apical part of P1, J1-2, L2 and P2 regions displayed lower sequence variability (Fig. 3a), and the substitution of these residues (mutants S1.2, S1.3 and S1.4) had a dramatic effect on the inhibition capacity of these constructs (Fig. 3b).

**Fig. 3.**
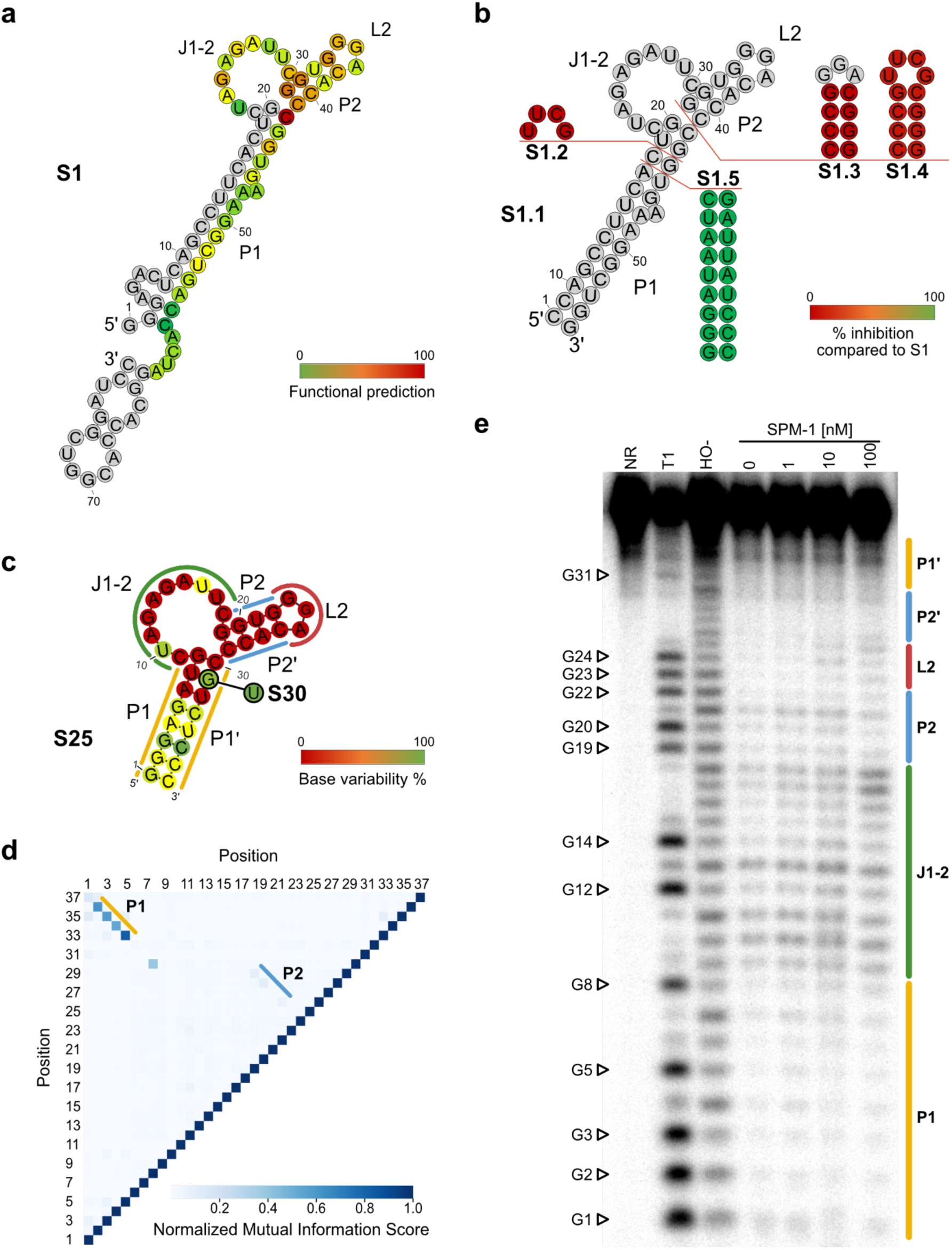
Secondary structure model of SPM-1-binding aptamers and sequence optimization. **a**, Secondary structure of S1 aptamer as predicted by RNA-fold. Stems (P), junction (J) and loops (L) are indicated. The functional contribution of each nucleotide to the function of clade 20 aptamers contained in F12.3 library was predicted by subtracting, for each nucleotide, the probability (expressed in bit) of finding this nucleotide in a sequence that was enriched from that of finding the same nucleotide in a sequence that was depleted. Summing the bit differences of each possible nucleotide at a given position and normalizing this value between 0 (completely counter selected nucleotide) and 100 (absolute conservation of the nucleotide) gave access to the functional prediction score that was color-coded. Note that constant regions were not considered in the calculations and were colored in gray. **b**, Impact of mutations and truncations on aptamer inhibition capacity. Different constructs were prepared and functionally evaluated. The inhibition capacity of each construct was normalized to that of S1 and color-coded. **c**, Predicted secondary structure of S25 aptamer and tolerance to sequence variability. The enriched pool D3.3 obtained at the end of doped-selection and functional screening was sequenced and sequence variation at each position was computed. Invariant nucleotides were easily discriminated from positions where variations were tolerated. Colored lines refer to the structures confirmed by in-line probing. The G to U mutation converting S25 into S30 is also indicated. **d**, Sequence covariations in the variants contained in D3.3 library. A covariation matrix was computed from enriched sequences contained in library D3.3 as described elsewhere^38^. Elevated Normalized Mutual Information score is indicative of possible base-pair formation and allows stem P1 and P2 to be identified. **e**, In-line probing of S25 performed in the presence or absence of SPM-1. A non-reacted control (NR), RNase T1 digest, alkaline-digest (HO^-^), a sample without SPM-1 (0) and three samples with SPM-1 (1, 10, 100 nM) are shown. G positions are marked with white arrows. Positions of structural elements of S25 are delineated by colored lines.

To refine aptamer characterization and possibly identify improved mutants, we next set-up a doped-SELEX experiment. The aptamer was first shortened down to 37 nucleotides by reducing the P1 helix down to 8 base pairs while preserving the rest of the molecule. This minimal S25 variant (Fig. 3c) conserved an overall affinity for SPM-1 and inhibition capacity nearing that of the full-length S1 aptamer (Supplementary Table 3). A mutant library was next prepared by randomizing each position to 9% prior to subjecting it to three rounds of SELEX followed by three rounds of µIVC screening (Supplementary Fig. 7a-d). Consistent with previous observations, sequence permutations preserving base pairing accumulated along P1 (Fig. 3c and Supplementary Fig. 7e-f), confirming that P1 primarily acts by providing stability to the molecule. Sequence covariation analysis of the selected molecules also highlighted the existence of P1 and P2 pairings, although P2 was less tolerant to sequence variation (Fig. 3d). A covariation was also detected between positions 8 and 30 although occurring at a much lower frequency (Fig. 3d). Interestingly, whereas the expected rigidity of paired regions was readily observable within P1 by in-line probing, P2 appeared to be more flexible (Fig. 3e), suggesting that this region could adopt several conformations (see below). Conversely to P1, the sequence of the rest of the molecule remained largely invariant except for positions 10, 31, and 16 to a lower extent (Fig. 3c and Supplementary Fig. 7e). Whereas position 16 was not further studied, U10A and G31U mutants were prepared and functionally evaluated (Supplementary Fig. 7f). Introducing the U10A mutation did not significantly affect the inhibition capacity of the aptamer whereas it severely impacted its transcription efficiency (Supplementary Fig. 7g). Elucidating the origin of this adverse effect would require additional investigations out of the scope of this study, leading us to disregard this mutant in subsequent analyses. G31U mutation converts a wobble pairing U7oG31 into a non-canonical U7oU31. Accordingly, these two base pairs were not only the most abundant species identified at the end of the process, but they were also the best tolerated among all the tested permutations (Supplementary Fig. 7g). In particular, introducing a G at position 7 or stabilizing this base-pair by adding an extra H-bond through the formation of G=C or C=G base-pairs (*i.e.*, S35 or S40 mutants) significantly impaired the function of the aptamer, suggesting that some flexibility should be preserved in this region as later confirmed using NMR (see below). Altogether, these data suggest that S25 and its G31U derivative (hereafter called S30) should be near-optimal SPM-1 inhibitors.

### Properties of the inhibitory aptamers

Evaluating the aptamer/SPM-1 complex formation using biolayer interferometry (BLI) revealed that the parental S1 aptamer and its two shorter derivatives (S25 and S30) shared similar affinities for SPM-1 with nM range (24 - 54 nM) *K_D_* values (Fig. 4a, Supplementary Fig. 8 and Supplementary Table 3). Interestingly, although backbone chemistry (RNA or 2’-FY) did not affect the *K_D_* value, we noticed that fluorinated aptamers associated slower with SPM-1 than their RNA counterparts. However, upon formation complexes involving 2’-FY aptamers also displayed slower dissociation rates. Electrophoretic mobility shift assay (EMSA) experiments confirmed aptamers/SPM-1 interactions with an even higher apparent affinity (Fig. 4b, Supplementary Fig. 9 and Supplementary Table 3), suggesting that, in BLI measurements, the aptamer-binding site might have been partly occulted upon SPM-1 immobilization. S25 specificity was confirmed by EMSA and beta-lactamase activity assay as target binding and inhibition was observed with SPM-1 but not with other tested beta-lactamases (Fig. 4c). Finally, we confirmed that the interaction readily depends on the sequence of the aptamer since a control with scrambled sequence (identical nucleotide compositions but in different order) did not display any measurable interaction (Fig. 4b, Supplementary Table 3).

**Fig. 4.**
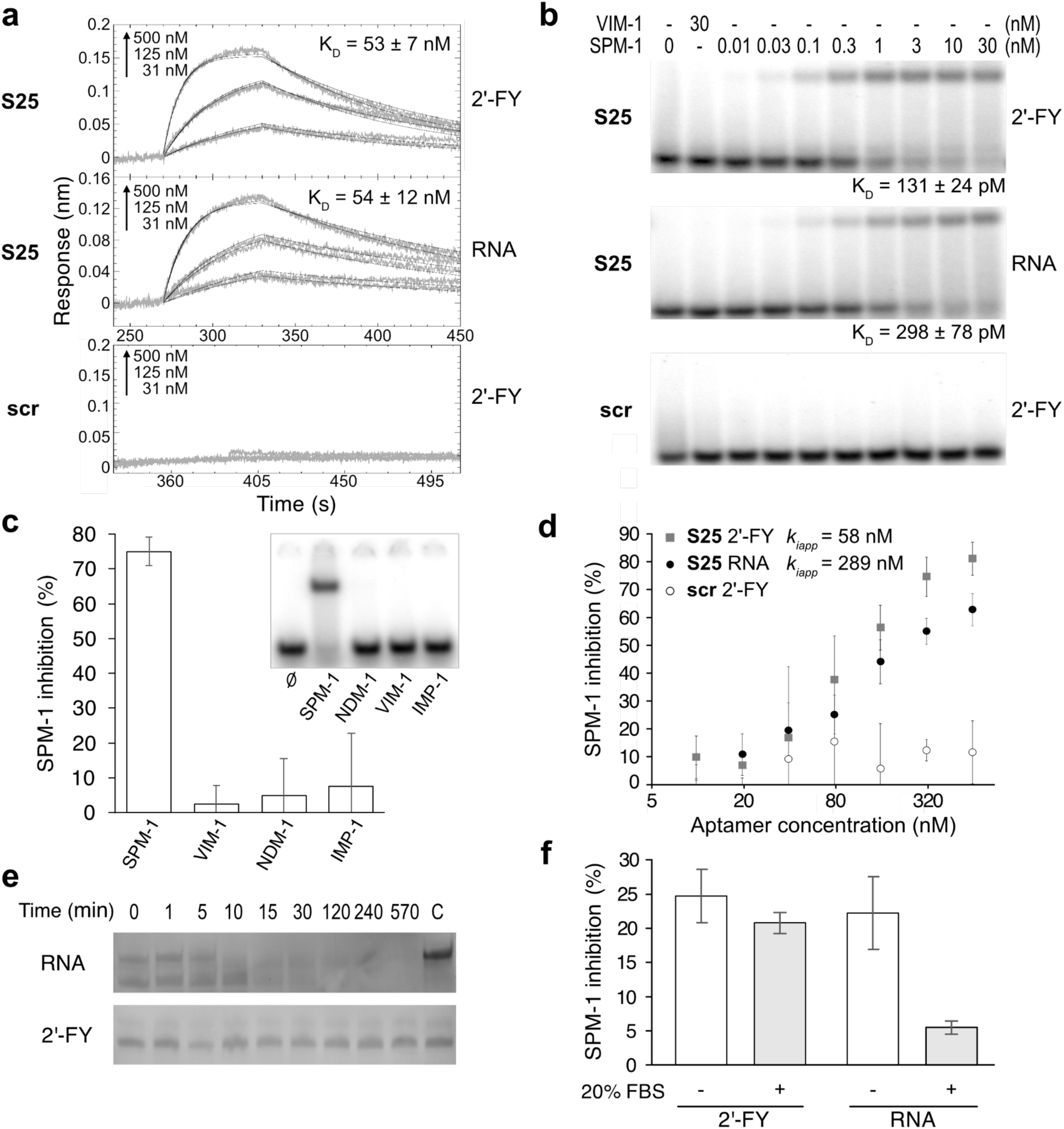
Characterization of the interaction between S25 inhibitory aptamer and SPM-1 beta-lactamase. **a**, Binding kinetics of S25 2’-FY and RNA monitored by BLI. **b**, Evaluation of dissociation constant between S25 and SPM-1 using electrophoretic mobility shift assay. Radiolabeled S25 was prepared in RNA or 2’-FY chemistry and incubated with a range of protein concentrations. VIM-1 and 2’-FY scrambled control (scr) were respectively used as protein and nucleic acid negative controls. *K_D_* values were then determined by fitting the data to an Hill equation. **c.** Specificity of binding and inhibition of S25 2’-FY. The aptamer was incubated with the target (SPM-1) or non-target beta-lactamases (NDM-1, VIM-1 and IMP-1) and its activity inhibition capacity as well as its binding capacity (inset) were evaluated. **d**. Effect of different concentrations of S25 2’-FY and RNA and scr 2’-FY on SPM-1 activity monitored using imipenem-based assay. **e**, Characterization of S25 aptamer nuclease resistance in a complex biological medium. S25 was prepared in RNA and 2’-FY chemistries prior to being incubated in 1% fetal bovine serum (FBS). Aptamer integrity was next evaluated by denaturing gel electrophoresis followed by ethidium bromide staining. **f**, Functionality of S25 aptamer in a complex biological medium. An excess of S25 RNA or 2’-FY was incubated with SPM-1 for 30 minutes in the presence or absence of 20% FBS. Beta-lactamase activity was then monitored using CCF2-FA assay.Values are the mean of 3 independent experiments and the error bars correspond to the standard deviation. Error bars were cut upon axis interception for an illustrative purpose.

We then tested the capacity of our aptamers to inhibit the SPM-1-mediated degradation of different beta-lactam substrates (Fig. 4d and Supplementary Table 3). As before, the shortening of the molecule had only a limited impact on its inhibition capacity with sub-micromolar to nanomolar apparent *k_i_* values. Interestingly, the backbone chemistry had a much higher effect since 2’-FY aptamers were 4 times more efficient inhibitors than their RNA counterparts, probably as the result of their lower dissociation from the target. SPM-1 inhibition was not only observed with the fluorescent substrate analogue (CCF2-FA) originally used to identify the aptamers, but also with the chromogenic CENTA and the unmodified imipenem (Supplementary Table 3), confirming the generic action of our inhibitors. Although S25 and S30 aptamers were identified in media supplemented with 5 mM MgCl_2_, maximal enzyme inhibition was reached at 2.5 and 1 mM MgCl_2_ respectively (Supplementary Fig. 10). Kinetic characterizations also suggested that the aptamers do not directly compete for substrate binding but rather behaved as uncompetitive inhibitors (Supplementary Fig. 11g).

We next evaluated the performances of S25 and S30 aptamers in more biologically relevant conditions. As expected, S30 RNA aptamer displayed a short <1 minute half-life in only 1% fetal bovine serum (FBS), whereas the integrity of 2’-FY aptamers was completely preserved (Fig. 4e and Supplementary Fig. 12). Modified aptamers remained even stable and functional for several hours in 20% FBS (Fig. 4f), demonstrating that, as expected, these inhibitors preserve their properties in a biological environment (*i.e.*, containing many other proteins, lipids, and RNAse activity) relevant to future applications.

### Structural rigidity of S25 and S30 aptamers

We further characterized our inhibitory aptamers using Nuclear Magnetic Resonance (NMR). Collected spectra display imino protons resonances between 9 and 15 ppm provided that they are involved in hydrogen bonds.

S25 RNA was first analyzed in Mg^2+^-free conditions at 15°C. The UoG wobble base pair was easily identified from the strong NOE between the G31-H1 and U7-H3 observable in the NOESY experiment at short mixing time (*i.e.*, 50 msec). The A-U Watson-Crick base pairs were next discriminated from G=C base pairs by the very strong correlation between U-H3 and A-H2. Besides, two strong NOEs are observable between G-H1 and the cytosine amino protons in a G=C pair, allowing base-paired imino protons in P1 and P2 stems to be assigned (Fig. 5a left and middle). Additional resonances and broad imino protons indicative of a weak stacking and interactions, arising from residues located outside the P1 and P2 stems could not be identified from analysis of 2D NOESY spectra. This underlines the dynamical behavior of the apical part of S25 RNA contrasting with the rigid P1 stem, supporting data from in-line probing experiments (see above). Although the aptamer was originally isolated in the presence of 5 mM MgCl_2_, NMR results indicate that the addition of this cation did not induce significant changes in the spectra. An accurate melting temperature (*Tm*) value could not be determined for this aptamer due to undefined lower and upper baselines of the melting curves, even in the absence of magnesium ions (Fig. 5a, right). However, as expected, the addition of the magnesium chloride increased the melting temperature (*Tm*) indicating an enhancement in the thermal stability of the aptamer. Interestingly, further increase of *Tm* values was observed upon 2’-FY backbone chemistry incorporation, demonstrating a stabilization of the molecule conferred by the modification.

**Fig. 5.**
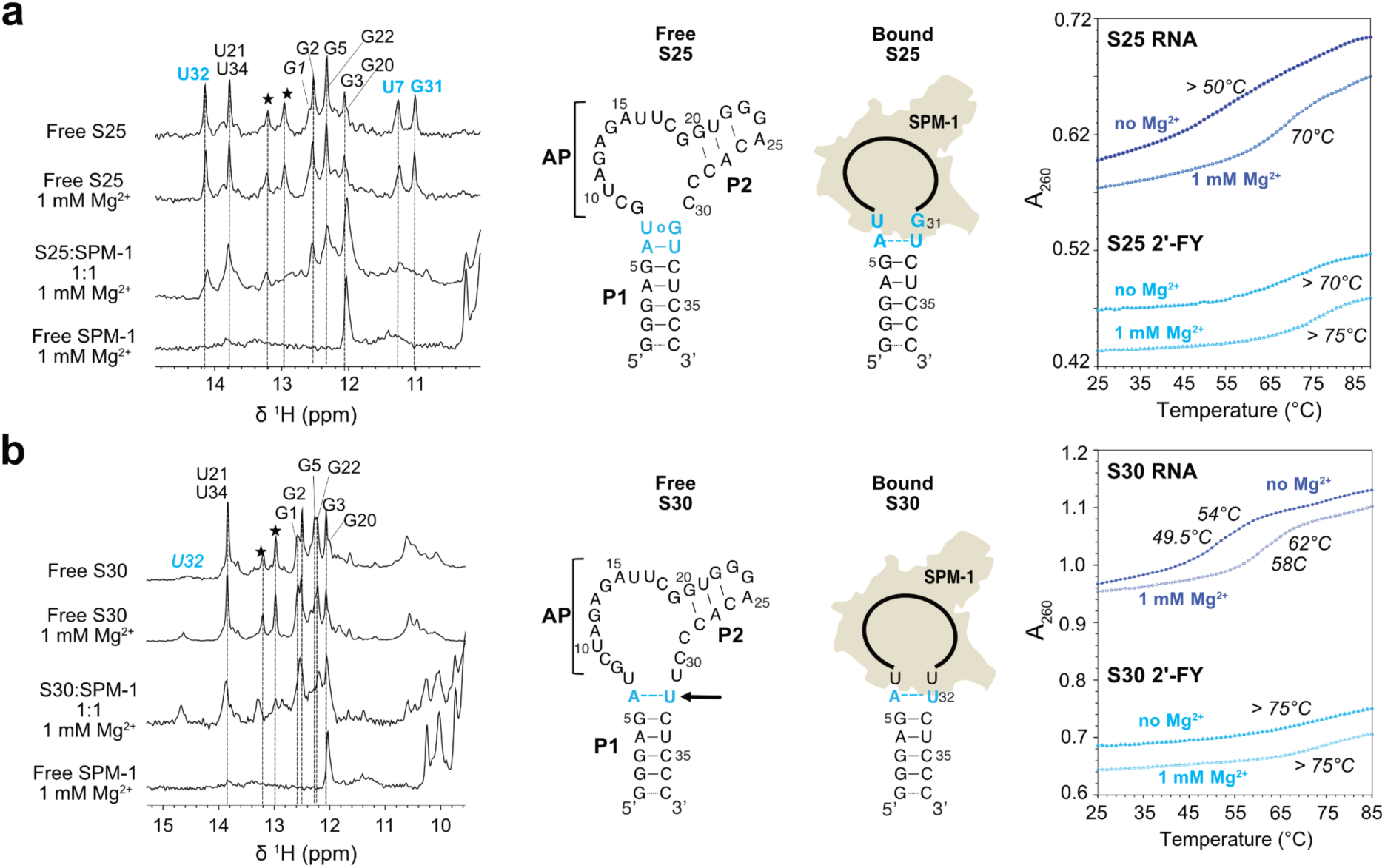
Solution secondary structures and stability of S25 and S30 RNA aptamers and their interaction with SPM-1 protein. **a**, S25 aptamer characterization. Left: Imino protons region of 1D NMR spectra recorded at 15°C in 50 mM sodium phosphate buffer (pH 6.4) in the absence of magnesium ions (Top), in the presence of 1 mM MgCl_2_ and for RNA:SPM-1 complex at a 1:1 ratio. The spectrum of the free protein recorded in the same condition is also shown (bottom). All imino protons were assigned via sequential NOEs observed in 2D NOESY experiments. Stars indicate resonances that could not be assigned. Middle: Secondary structure models of free and bound S25. Dashed lines indicate weak hydrogen bonds. “AP” indicates the apical part of the aptamer which is defined from residues G8 to C30. Right: UV melting curves at 260 nm absorbance in the absence of magnesium ions and at 1 mM MgCl_2_ for S25 RNA (dark blue) and 2’-FY (light blue). **b**, S30 aptamer characterization. Left: Imino protons region of 1D NMR spectra recorded at 15°C in 50 mM sodium phosphate buffer (pH 6.4) in the absence of magnesium (Top), in the presence of 1 mM MgCl_2_ and for RNA:SPM-1 complex at a 1:1 ratio. The spectrum of the free protein recorded in the same condition is shown (bottom). All imino protons were assigned through sequential NOEs observed in 2D NOESY experiments. Stars indicate resonances that could not be assigned. Middle: Secondary structure models of free and bound S30. Dashed lines indicate weak hydrogen bonds. The arrow indicates the A-U base-pair stabilized upon magnesium addition. “AP” indicates the apical part of the aptamer which is defined from residues G8 to C30. Right: UV melting curves at 260 nm absorbance in the absence of magnesium and at 1 mM MgCl_2_ for S30 RNA (dark blue) and 2’-FY (light blue).

S30 RNA imino protons were also assigned using U34-H3 as a starting point. As observed for S25, NMR analysis revealed that the P1 stem is formed although A6-U32 involved a weak H-bond and the predicted U7oU31 was not detectable (Fig. 5b, left). Moreover, similarly to S25, the apical part of the aptamer remained flexible as indicated by weak sequential NOE observed in the P2 stem. Yet, a slight stabilization of the A6-U32 pair was obtained upon addition of magnesium chloride. The lower overall stability of S30 was also apparent on melting curves whose derivative displayed two close transitions at 49.5 and 54.0°C. As for S25, an increased *Tm* was observed upon addition of magnesium chloride (Fig. 5b, right). Interestingly, preparing S30 in 2’-FY chemistry endowed the molecule with a thermal stability similar to that of S25 further highlighting the benefit brought by the incorporation of fluorinated nucleotides, especially at the level of this U_7_oU_31_ base pair.

We next used NMR to gain some insights on the interaction between SPM-1 protein and the aptamers. The chemical shifts of imino protons were monitored as a function of the protein concentration to map the RNA regions in contact with SPM-1. Successive addition of the enzyme to S25 and S30 resulted in several spectral changes mostly involving the apical part of the RNA as illustrated by the broadening and disappearance of the corresponding resonances (Fig. 5a, left and 5b, left). Interestingly, besides these effects, in S25, the G31 and U7 resonances disappeared upon protein binding, revealing modifications of solvent accessibility. As the exchange rate of imino proton with the solvent is related to the dissociation constant of a base pair, this observation can be interpreted as a melting of the U7oG31 base pair. Concomitantly, the U32 resonance broadened, indicating a destabilization of the A6-U32 base pair upon SPM-1 binding. By contrast, in S30, the P1 stem remained unaffected. Altogether, these data suggest that a reshaping of the aptamers apical part may occur upon protein binding. Yet, it cannot also be excluded that bound aptamers adopt conformations not detectable by NMR. Further investigations will be required to establish the detailed mechanism of binding of SPM-1 to S25 and S30 aptamers.

## Discussion

In this work, we introduced an ultrahigh-throughput aptamer discovery pipeline combining the use of *in vitro* selection in tandem with microfluidic-assisted functional screening, high-throughput sequencing and bioinformatics to rapidly identify potent inhibitors functional in challenging biological environments. We demonstrated the efficiency of this approach by discovering a modified aptamer able to efficiently and specifically inhibit SPM-1, a metallo-beta-lactamase particularly difficult to target for drug development^32^. Starting from a complex random library we first enriched the pool in SPM-1 binders prior to enriching it in efficient inhibitors. Moreover, whereas the first rounds were performed with libraries expressed as RNA, we rapidly switched to a mixed chemistry made of purines ribonucleotides and 2’-fluorinated deoxyribose pyrimidines in order to generate nuclease-resistant oligonucleotides compatible with future applications. Sequence analysis enabled the whole selection process to be characterized at once and variants of interest to be readily identified, avoiding long and tedious post-selection characterizations. Upon functional screening of SELEX-enriched libraries, clade 20 was identified as the dominant species, before deeper bioinformatics identified S1 aptamer as being the most promising candidate. Interestingly, most of the tested candidates displayed a capacity to inhibit SPM-1, confirming the efficiency of the process.

S1 was next successfully miniaturized down to the 37 nucleotide-long S25 aptamer. Further attempts to improve this aptamer revealed that it was likely an optimal sequence possessing all the features searched for such a molecule. First, the aptamer displayed high affinity and strong specificity for its target. Second, this interaction led to a very strong inhibition of SPM-1 activity and prevented the transformation of every tested antibiotic and substrate analogues. Third, the small size of the aptamer makes it producible both by solid phase and enzymatic synthesis. Last but not least, the use of 2’-FY chemistry not only provided the expected nuclease-resistance to the molecule but it also endowed it with a superior structural rigidity dropping the need of magnesium ions to the low mM concentration typically found in biological environments. Although the proper understanding of the molecular recognition between the aptamers and their target protein will require deeper characterizations, NMR data suggest that the P1 stem provides a rigid scaffold supporting the apical region of the aptamer involved in protein recognition. The upper part encompassing the weak P2 stem displays higher flexibility and collected data suggest that recognition can take place through an induced fit mechanism. Inhibition kinetics also suggest that S25 may primarily work as an uncompetitive inhibitor, a scenario previously described for another metallo-beta-lactamase targeting aptamer^39^.

Performing functional screenings using droplet microfluidics was instrumental in two ways. First, this technology is unique in allowing the parallel analysis of million variants while physically confining the genotype with its encoded phenotype, a feature particularly relevant with diffusive phenotypes as it is often the case with the enzymatic substrate conversion. Whereas other screening formats exploiting molecular capture at the surface of beads or at the surface of sequencing flow-cells have been described^40^, none of them would enable to preserve together the tested aptamer, the target enzyme together with reaction products. Second, the strong miniaturization of the reaction vessels enables precious reagents to be used at low cost and allows millions of variants to be functionally screened while being expressed as modified aptamers. This contrasts with two-step strategies that have been frequently used and in which hits are first selected in cheaper RNA or DNA chemistry prior to modifying the backbone of the best candidates. Although attractive on an economical point of view, such a strategy carries the risk of developing sub-optimal molecules with some positions intolerant to modification as illustrated by the development of Macugen^17^. Indeed, the parental aptamer of this therapeutics was first developed in RNA chemistry and the backbone of the best hits was later modified. Whereas pyrimidines 2’-fluorination was somehow well tolerated, two purines remained intolerant to 2’-O-Methylation, leaving the aptamer only partly modified, so protected^41^. Interestingly, a similar scenario could have happened in the present work as witnessed by the adaptation of the library composition upon change in backbone chemistry. Consistently, enriched libraries expressed as RNA performed better in this chemistry than in 2’-FY, whereas mirror situation was observed when enriched libraries were initially expressed as 2’-FY oligonucleotides. These observations highlight the importance of introducing modified nucleotides as early as possible in the discovery pipeline. In this work, we chose to switch the pool chemistry in the course of the study to evaluate its impact on pool content. Nevertheless, it makes no doubt that the process could begin with a starting library already expressed as a modified pool. 2’ fluorination of pyrimidine is a modification typically used in several drugs, but new triphosphate modified nucleotides are regularly introduced together with polymerases allowing their incorporation^33^. Combining the use of these new chemistries with the pipeline disclosed in the present work could pave the way toward the efficient discovery of a new generation of drugs in the form of potent inhibitors against enzymes difficult to target (like the metallo-beta-lactamase used in this study) or even considered as undruggable^42^.

## Online content

Any methods, source data, supplementary information, acknowledgements, peer review information; details of author contributions and competing interests; and statements of data and code availability are available at http://doi. …

## Online content

## Methods

### General information

Chemicals were purchased from Sigma-Aldrich and used without further purification unless otherwise stated. Molecular biology enzymes were purchased from New England Biolabs unless otherwise stated. Unless explicitly mentioned, DNA oligonucleotides were obtained from Integrated DNA Technologies.

### Protein preparation

The plasmid SPM1-pET24a+ containing the gene encoding SPM-1 (N-terminal His-tagged, derived from *Pseudomonas aeruginosa*) under the control of a T7 RNA polymerase promoter was transformed into *Escherichia coli* strain BL21-DE3(NEB). After culture and induction in the presence of kanamycin, 400 µM IPTG and 100 µM ZnCl_2_, the protein was purified by nickel affinity chromatography and gel filtration (Superdex 200). The protein was then stored at -20°C in a conservation buffer (50 mM TRIS-HCl pH 8.0, 500 mM NaCl, 5% (v/v) Glycerol and 50 µM ZnCl_2_).

### RNA and 2’-FY RNA preparation

RNA was transcribed from PCR-amplified templates. For this, two oligonucleotides were designed with an overlap of 30 nucleotides and amplified using Q5 High-Fidelity DNA polymerase (NEB) according to the supplier’s instructions. After ethanol precipitation, the DNA template was used for *in vitro* transcription with T7 RNA polymerase (homemade) as reported previously^43^. Aptamers in 2’FY were transcribed with the DuraScribe T7 transcription kit (Biosearch Technologies) according to the supplier’s instructions. Aptamers were gel-purified, and their concentration was determined by spectrophotometry (NanoDrop 1000 Spectrophotometer, Thermo Scientific).

### SELEX

The initial RNA pool used for SELEX consisted of a 40 nucleotides randomized region flanked by constant regions (5’constant: 5’-GGA GCT CAG CCT TCA CTG-3’, 3’constant: 5’-GGC ACC ACG GTC GGA TCC-3’, Supplementary Fig. 1)^44^. After each round, the pool was recovered via RT-PCR using a primer annealing to the 3’ constant region, while a primer binding in the 5’ constant region added the T7 promoter necessary for *in vitro* transcription (fwd_primer: 5’-AAT TCT AAT ACG ACT CAC TAT AGG AGC TCA GCC TTC ACT GC-3’; rev_primer: 5’-GGA TCC GAC CGT GGT GCC-3’). The DNA pool was obtained by PCR using Q5 DNA polymerase (New England Biolabs) amplifying a reverse complementary oligonucleotide including the randomized region and the 3’ constant region with the primer fwd_primer (Supplementary Table 2). For the first round of selection, 800 pmoles of DNA template were *in vitro* transcribed using T7 RNA polymerase (homemade). From round 7 on, the pool was transcribed in 2’-FY using the DuraScribe T7 transcription kit (Biosearch Technologies). For every round *α*-^32^P-ATP (Hartmann Analytic) was added to the *in vitro* transcription for radioactive body-labeling of the RNA. After transcription, products were precipitated, resuspended in ddH_2_O, and the concentration was determined with a liquid scintillation analyzer (Tri-Carb 2800TR, Perkin Elmer). For correct folding, the RNA was heated up to 95°C for 5 min and snap-cooled on ice for 5 min. 2 nmoles of RNA were mixed with 5 µg yeast tRNA (Invitrogen) and SELEX buffer (SB: 20 mM TRIS-HCl pH 7.4, 100 mM KOAc, 5 mM MgCl_2_, 5 µM ZnCl_2_, 0.01% Tween-20). For selection the His-tagged target protein SPM-1 was immobilized to Co^2+^-NTA magnetic beads (Dynabeads, Thermo Fisher Scientific). For protein immobilization, 500 pmoles SPM-1 were incubated in SB on the beads rotating for 15 min at room temperature. The beads were collected with a magnetic separation rack, the supernatant discarded, and the beads washed three times with SB. Also, beads without SPM-1 were prepared in SB and used for pre-selection and measurement of bead-binding RNAs. For pre-selection, the RNA pool was incubated with uncoupled beads rotating for 15 min at 37°C to remove any RNA that bound to the beads. The unbound RNAs in solution were separated from the beads and transferred to a new tube with either beads coupled with SPM-1 for subsequent selection or beads without SPM-1 to measure binding of the RNA pool to the beads. The RNA pool was incubated with the target protein for 30 min at 37°C under rotation. Afterwards, the beads were collected, the supernatant discarded, and the beads washed with SB. To recover the protein from the beads with their bound RNA, an elution buffer (SB with 300 mM imidazole) was added to the beads and the beads were left rotating at room temperature for 5 min. The solution was separated from the beads, transferred to a new tube and the RNA was recovered by phenol-chloroform extraction. The fraction of RNA bound to the protein or to the beads were measured with a liquid scintillation analyzer and calculated as percentage of the input. The recovered RNA was then reverse transcribed using SuperScriptII (Thermo Fisher Scientific) and PCR amplified with Taq DNA polymerase (homebrew) in a one-step setup. The cDNA pool was then stored and used for *in vitro* transcription of the RNA pool for the next selection round. For doped SELEX, the selection was performed as described above with TALON beads (Takara Bio), 2.5 nmoles RNA and 250 pmoles of SPM-1 in selection.

### Microfluidic Inhibition Screening

*Library preparation:* The starting library of DNA templates recovered after the SELEX procedure was PCR amplified in 100 µL of SsoFast EvaGreen Supermix (Bio-Rad) supplemented with 20 pmoles of Ampli-Forward (designed to introduce the T7 RNA polymerase promoter and a modified ITS known to increase the transcription yield) and 20 pmoles of Ampli-Reverse primers (Supplementary Table 2). PCR products were used as a template in a second reaction using the same reaction mixture but using as primers 20 pmoles Add-barcode primer (insert 20 randomized nucleotides acting as Unique Droplet Identifier, UDI, upstream the construct)^26^, and 20 pmoles Ampli-Reverse primer. PCR products were finally purified using the SPRIselect Bead (Beackman Coulter).

*Microfluidics-assisted functional screening:* Microfluidic chips were prepared as described before^26^, and the three main steps of the screening were performed as follows:

i. Digital droplet PCR: a dilution of purified DNA (calculated to reach the desired droplet occupancy) was prepared in 200 µg/mL total yeast RNA (Ambion) before introduction into a PCR medium containing SsoFast EvaGreen Supermix (Bio-Rad) at the recommended concentration, 0.1% Pluronic F68 (Sigma), 0.2 μM of each primer (Ampli-Barcode and Ampli-Reverse), and 1 μM of Cyanine 5 (Cy5; red-fluorescence droplet tracer). The resulting solution was injected into a droplet generator device and dispersed into 2.5 pL droplets carried by a Novec 7500 fluorinated oil phase (3M) containing 3% surfactant to stabilize the emulsion. The emulsion was collected and thermocycled to allow DNA amplification.
ii. Droplet fusion: addition of *in vitro* transcription medium and RNA production. An *in vitro* transcription medium (IVT) containing 4 mM of each rNTP (Larova) or 2’-fluoro-dCTP and 2’-fluoro-dUTP (Jena Bioscience), 44 mM TRIS-HCl pH 8.1 at 37°C, 22 mM MgCl_2_, 500 µM DTT, 1 mM spermidine, 1 µg inorganic pyrophosphatase (Roche), 17.5 μg/mL T7 mutant K378R, Y639F, H784A^45^ bacteriophage RNA polymerase (produced in the laboratory), 5% DMSO, 0.1% Triton X100, 0.1% Pluronic F68, 0.1 µM Cy5, and 7 nM purified SPM1. The mixture was injected into a droplet fusion device and dispersed in 17.5 pL droplets carried by Novec 7500 oil phase supplemented with 2% surfactant. Droplets containing PCR-amplified DNA were re-injected into the chip through another inlet and synchronized one-to-one with IVT droplets prior to fusing these pairs by applying an AC electric field (700 V, 70 kHz). The emulsion was collected at 4°C and then placed for 2 h at 37°C.
iii. Droplets injection/incubation/sorting and recovery of genes of interest. Droplets were reinjected into an integrated device (Supplementary Fig. 2) mounted onto a thermoplate (Tokai) set at 42°C to favor the isolation of stable aptamers and spaced with a surfactant-free Novec 7500 oil phase. Around 3 pL of a mixture of CCF2-FA (Thermo-Fisher) diluted in 50% DMSO, 20 mM TRIS-HCl pH 8.1 at 37°C and 5 µM ZnCl_2_ were piconinjected into each droplet. After 5-10 min incubation in the delay line, red (Cy5, droplet tracer), blue and green (CCF2-FA emissions) fluorescence of each droplet was analyzed and used to sort the droplets of interest by applying an AC electric field (1200 V and 30 kHz) to deflect droplets of interest into the “sort” channel. The recovered emulsion was then broken with 100 μL of 1H,1H,2H,2H-Perfluoro-1-octanol (Sigma-Aldrich), and the aqueous phase was mixed with 100 μL of 200 μg/μL total yeast RNA (Ambion).

### Enrichment evaluation and regeneration of the libraries

Recovered DNA libraries were PCR-amplified as above using Ampli-Forward and Ampli-Reverse primers. PCR products were then *in vitro* transcribed in the presence of 7 nM SPM-1 for 1h at 37°C. Then a mixture of CCF2-FA (Thermo-Fisher) prepared in 50% DMSO, 20 mM TRIS-HCl pH 8.1 at 37°C and zinc chloride was added to obtain a final concentration of 2 µM CCF2-FA and 5 µM ZnCl_2_. CCF2-FA degradation was monitored by recording the fluorescence emitted at 450 nm and 520 nm upon excitation at 375 nm every minute on a microplate reader (SpectraMax iD3, Molecular Devices). Upon each round of screening, UDI barcodes were reset to preserve their uniqueness. To do so, 1 µL of the DNA template contained in the recovered aqueous phase was diluted in 100 µL of SsoFast EvaGreen Supermix (Bio-Rad) PCR mixture prior to reamplifying it by PCR using 20 pM Add-barcode and 20 pM Ampli-Reverse primers (Supplementary Table 2). PCR products were purified using the SPRIselect Bead (Beackman Coulter) and quantified using spectrophotometry (UV-vis spectrophotometer DS-8X).

### Sequence analysis of the enriched libraries

After each validated round of screening, DNA recovered from the sorted droplets was amplified using Ampli-Barcode and NGS-Reverse primers (Supplementary Table 2) to append P5 and P7 sequences subsequently used for molecular indexing using the Nextera Indexing kit (Illumina) as recommended by the supplier. PCR products were purified using the SPRIselect system (Beckman Coulter) prior to quality check by microcapillary electrophoresis (Bioanalyzer, Agilent). Following quantification by Qubit (Thermo Scientific), obtained libraries were analyzed on a V3-150 chip (Illumina) by MiSeq sequencing device (Illumina).

### NGS data analysis

Raw NGS data were analyzed using a custom python workflow. Reads with a Phred quality score below 30 and those that fell below a manually set occurrence threshold were excluded from further analysis. Data treatment and results visualization were performed using Python libraries including Pandas^46^ for data manipulation and analysis, Matplotlib^47^ and Seaborn^48^ for plot generation.

To extract the unique droplet identifier UDI^26^ and the 40 randomized nucleotides, the regular expression ^AA(.{20}.) and (.{40})GGCACCACGGTCGGATCC were employed respectively. In the case of doped NGS data, the mutated sequences were extracted using CTTCACTGC(.*)GGCACCACGGTCGGATCC. After removing repeated sequences (sharing the same UDI and sequence) each UDI and its corresponding sequence were row organized into a Pandas data frame. Reads displaying the same sequence but different barcodes (coming from different droplets) were summed and used to calculate the sequence occurrence and frequency in each library at each selection round.

To identify the enriched motifs through the selection process, a FASTA file was generated with the filtered randomized sequences from each selection round. This FASTA file was then analyzed using the MEME suite^49^. The MEME algorithm was used to identify up to 10 significant enriched motifs per selection round. The FIMO algorithm was then used to assign to each sequence its corresponding motif. Because the same motif can evolve through the selection process, all the identified motifs of all selection rounds were clustered in clades according to its phylogenetic similarity. Briefly, Clustal Omega algorithm^50^ was used to align all the motifs, then the Biopython’s^51^ Bio.phylo^52^ package was used to calculate the distance between the aligned sequences. Motifs presenting a distance less than 0.3 units in the distance matrix were clustered together in clades. This clustering revealed distinct groups of motifs, which were then visualized as clades for the different selection rounds using the NetworkX^53^ library. Finally, phylogenetic trees of the different clades were visualized using the Toytree^54^ library. The most enriched clade (clade 20) was then analyzed at the single sequence level. Upon sequence filtering according to their cumulated frequencies > 0.1% from rounds F12 to F12.3, sequences were organized according to their enrichment slope (calculated according to the sequence frequency in each selection round) and represented in the forms of a row normalized HeatMap of rounds F12 to F12.3 and with the logarithm of the frequency of the round F12.3.

To investigate the covariation between nucleotide positions, we calculated the normalized mutual information (NMI)^38^ score for each pair of positions. We used the normalized_mutual_info_score function from scikit-learn^55^ to calculate the NMI score for each pair of positions (i, j). This resulted in a matrix of NMI scores, where the score at position (i, j) represents the strength of covariation between positions i and j. The matrix was visualized as a heatmap using the seaborn library, with the x-axis representing the nucleotide positions and the y-axis representing the same positions in reverse order. The color bar indicates the magnitude of the NMI score, ranging from 0 (no covariation) to 1 (perfect covariation).

Venn diagrams were generated using the Matplotlib-Venn library with the unique sequences at the selection rounds. Logos of the doped selection were made using the WebLogo API^56^.

### In-line probing

For in-line probing, RNA was dephosphorylated and ^32^P-labeled at the 5’ end as previously described^43^. The labeled RNA was gel-purified and 35 kCPM of the purified ^32^P-labeled RNA were incubated for 68 h at 22°C in in-line reaction buffer (10 mM TRIS-HCl pH 8.3, 10 mM MgCl_2_, 100 mM KCl) with and without protein. To generate a size marker, the ^32^P-labeled RNAs were subjected to alkaline hydrolysis by incubation for 3 min at 96°C in 50 mM Na_2_CO_3_ pH 9.0 or incubated for 3 min at 55°C with 20 U RNase T1 at denaturing conditions to identify guanine residues. After in-line reaction, all reactions were ethanol-precipitated, and the pellets were dissolved in 5 M urea. All reactions were separated by denaturing polyacrylamide gel electrophoresis. Afterward, gels were dried and analyzed using phosphoimaging (Amersham Typhoon, GE Healthcare).

### Electrophoretic mobility shift assay

To determine the binding affinity of the aptamers to SPM-1, electrophoretic mobility shift assay (EMSA) was performed. After purification by preparative polyacrylamide gel, the aptamers were dephosphorylated with calf intestinal phosphatase (Roche) and radioactive end-labeled using *γ*-^32^P-ATP and T4 polynucleotide kinase (Roche) according to the manufacturer. Unincorporated *γ*-^32^P-ATP was removed using G25 microspin columns (Cytiva). The radioactive activity of the aptamer was measured, adjusted for each reaction to 10,000 cpm of radioactive labeled aptamer in SB. The prepared aptamer was mixed with different amounts of SPM-1, with final concentrations ranging from 10 pM to 30 nM. The samples were incubated at 37°C for 30 min. After incubation the samples were immediately loaded onto a 6% native PAA gel and run at 100 V at 4°C in TBM (0.89 M TRIS-HCl pH 7.4, 0.89 M boric acid, 1.5 mM MgCl_2_). After running, the gel was dried for 30 min at 80°C and the radioactive signal of the aptamer was detected with a Typhoon phosphoimager (Cytiva). Signal intensity of the bound fraction in each reaction was measured with imageJ and a least square fit was calculated to model the binding dynamic based on the Hill equation to obtain the *K_D_* of the aptamer.

### BLI assay

Kinetic experiments were performed at 25°C on an Octet R8 instrument (Sartorius France S.A.S, Dourdan, France). The His-Tagged SMP-1 protein and the RNA molecules were prepared in 20 mM HEPES pH 7.5, 100 mM KOAc, 5 mM MgCl_2_, 5 µM ZnCl_2_ and 0.01% Tween-20 (running buffer). The protein was captured at 50 nM on Ni-NTA probes (Sartorius) for 90 s after a 60 s activation step with 5 mM NiCl_2_ prepared in mQ water. Under these conditions, 1.5-2.5 nm responses of captured targets were obtained. The association and dissociation phases were monitored for 60 and 120 s, respectively. One well of the 96-well plate filled with a running buffer was used to generate the blank sensorgram. The regeneration of the biosensor was achieved using 10 mM glycine pH 1.7 as recommended by the supplier. The obtained sensorgrams were analyzed using the Octet Analysis Studio 13.0.2.43 software (Sartorius) and fitted to a 1:1 model of interaction. The dissociation equilibrium constant, *K_D_*, was calculated as the ratio of the dissociation rate over the association rate constants. *K_D_*s are the average and standard deviation of at least two independent experiments with kinetics performed in duplicate (functionalized probes plunged twice in the same wells).

### Inhibition assays

Different concentrations (10 nM - 2 µM) of purified aptamer candidates were incubated in 20 mM HEPES pH 7.5, 100 mM KOAc, 5 mM MgCl_2_, 5 µM ZnCl_2_, 0.01% Tween-20, 1 mg/mL BSA for 10 min in the presence of SPM-1. Depending on the assay, SPM-1 activity was then monitored upon addition of different concentrations of substrates.

Imipenem assay: 800 µM of imipenem (Sigma) were added to 14 nM SPM-1 and substrate conversion was monitored at 37°C by continuously recording the absorbance at 297 nm.

CCF2-FA assay: 20 µM of CCF2-FA were added to 7 nM SPM-1 and substrate conversion was monitored at 37°C by recording the fluorescence emitted at 450 nm and 520 nm upon excitation at 375 nm.

CENTA assay: 100 µM CENTA (Sigma) were added to 7 nM SPM-1 and substrate conversion was monitored at 37°C by recording absorbance at 409 nm. Each measurement is repeated 2 or 3 times independently.

The substrate consumption rate was then computed at each inhibitor concentration and the values were plotted as a function of inhibitor concentration. The slope of 1/V (where V is the reaction rate) was then plotted as a function of inhibitor concentration to determine the *ki_app_* as the intercept of x-axis.

### Magnesium dependency

Upon purification, aptamer candidates were incubated at different concentration in 20 mM HEPES pH 7.5, 100 mM KCl, 5 µM ZnCl_2_, 0.01% Tween-20, 1 mg/mL BSA, and 14 nM SPM-1 for 10 min with different concentrations of magnesium chloride. SPM-1 activity was then monitored using imipenem degradation essay. The *ki_app_* was then computed for each magnesium concentration. Each measurement is repeated twice independently.

### Stability essay

Purified aptamers were incubated in 1% or 20% Fetal Bovine Serum (FBS) with 20 mM HEPES pH 7.5, 100 mM KOAc, 5 mM MgCl_2_, 5 µM ZnCl_2_, 0.01% Tween-20, at 37°C. Aptamer integrity was then verified by migration in a 12% acrylamide denaturing (8 M urea) gel.

200 nM of aptamer was incubated with 7 nM SPM-1 in 20 mM HEPES pH 7.5, 100 mM KOAc, 5 mM MgCl_2_, 5 µM ZnCl_2_, 0.01% Tween-20 at 37°C with or without 20% FBS. SPM-1 activity was then monitored usingCCF2-FA degradation assay.

### NMR experiments

NMR spectra were recorded at 700 MHz on an Avance III Bruker spectrometer equipped with a z-gradient TCI cryoprobe. NMR data were processed using TopSpin (Bruker) and analyzed with NMRFAM-SPARKY software packages^57^.

Free unlabeled S25 and S30 RNA samples volumes were 150 µL in 3 mm NMR tubes at concentrations ranging from 0.1 to 0.4 mM. NMR experiments were performed in 50 mM sodium phosphate at pH 6.4 and 150 mM NaCl in 90/10 H_2_O/D_2_O. One dimensional spectra were acquired at 10°C, 15°C, 20°C, 25°C, 30°C, 37°C and 45°C. Magnesium was finally added at a concentration of 1 mM. Solvent suppression was achieved using combined “Jump and Return” and WATERGATE sequences^58–60^. Two-dimensional NOESY spectra were acquired with 50 ms and 400 ms at 15°C. Base-pairing was established via sequential nuclear Overhauser effects (nOes) observed in 2D NOESY spectra at different mixing times.

RNA:SPM-1 complexes were formed by successive addition of the protein to the RNA by monitoring the imino protons region at 15°C in 50 mM sodium phosphate at pH 6.4, 150 mM NaCl and 1 mM MgCl_2_. Saturation of the RNA was observed at a 1:1 stoichiometry.

### UV melting experiments

Thermal denaturation of S25 and S30 aptamers were monitored on a CARY3500 UV/vis spectrophotometer (Agilent) equipped with an 8-position sample holder and a Peltier temperature control accessory. The experiments were performed at 1 µM in 50 mM sodium phosphate buffer, at pH 6.4, in 100 µL micro quartz cuvettes. The RNAs were refolded as described above and magnesium chloride was next added at a concentration of 1 mM. A cuvette that contained the buffer with no magnesium was used as a reference. Samples were overlaid with 100 µL of mineral oil to prevent evaporation at high temperature. An initial 10 min equilibrium time at 25°C was included prior to the temperature ramping. Denaturation of the samples was achieved by increasing the temperature at 1°C.min^-1^ from 25 to 90°C and monitored at 260 nm. The melting temperature (*Tm*) was determined as the maximum of the first derivative of the UV melting curves. Each experiment was repeated independently two or three times.

## Acknowledgements

The authors thank Anne-Caroline Jousset for her assistance with sequencing library validation as well as Sandrine Koechler and Abdelmalek Alioua (IBMP Gene Expression Analysis facility) for technical assistance with high-throughput sequencing. This research was funded by a joint Agence Nationale de la Recherche/Deutsche ForschungsGemeinschaft grant (project “DIRA”, ANR-19-CE44-0020) and Fondation pour la Recherche Médicale (PhD grant extension funding to CH). Performed within Interdisciplinary Thematic Institute “IMCBio, as part of the ITI 2021–2028 program of the University of Strasbourg, CNRS and Inserm, this work was supported by IdEx Unistra (ANR-10-IDEX-0002) and by SFRI-STRAT’US project and EUR IMCBio (ANR-17-EURE-0023) under the framework of the French Investments for the Future Program. We also thank structural Biophysico-Chemistry platform of the European Institute of Chemistry and Biology (IECB, UAR 3033 - US001) for the use of the BLI instrument, which was acquired in part with the financial support of La Région Nouvelle-Aquitaine, Surphase (San Sebastian, Spain), DIPC (San Sebastian, Spain), Novaptech (Gradignan, France) and CovalX Analytics SAS (Martillac, France). CD is grateful to the INSERM U1212 - CNRS UMR 5320 ARNA laboratory and the Euskampus foundation through the LTC Sarea program for financial support. This work used the Integrated Structural Biology platform of the Strasbourg Instruct-ERIC center IGBMC-CBI supported by FRISBI (ANR-10-INBS-0005). This work was also supported by the “Centre National de la Recherche Scientifique” (CNRS) and the “Université de Strasbourg”, from whom it received support from its Initiative of Excellence (IdEx).

## Contributions

B.S. and M.R. conceptualized the study, provided supervision, discussed the results and acquired funding. M.R. wrote the manuscript. C.H. and J.H. performed the experimental investigation, analyzed data and created Figures and Tables. R.C. was responsible for the bioinformatics analyses and created Figures and Tables. I.L. performed and analyzed the NMR experiments and created Figures, BK: provided support for NMR experiments, L.K. performed and analyzed the in-line probing experiments, E.P. prepared and analyzed the SPM-1 protein, S. B. prepared and optimized molds for microfluidic device fabrication and provided support with microfluidic optical set-up, C.D. performed and analyzed the BLI assays. All authors reviewed the manuscript and agreed on its content.

## Corresponding authors

Correspondence to

## References

1. Fersht, A. Enzyme Structure and Mechanism. (Freeman, New York, 1995).

2. Shapira, A. & Benhar, I. Toxin-Based Therapeutic Approaches. Toxins 2, 2519–2583 (2010).

3. Wencewicz, T. A. Crossroads of Antibiotic Resistance and Biosynthesis. J. Mol. Biol. 431, 3370–3399 (2019).

4. Kalera, K., Stothard, A. I., Woodruff, P. J. & Swarts, B. M. The role of chemoenzymatic synthesis in advancing trehalose analogues as tools for combatting bacterial pathogens. Chem. Commun. 56, 11528–11547 (2020).

5. Walsh, C. Suicide substrates: mechanism-based enzyme inactivators. Tetrahedron 38, 871– 909 (1982).

6. Chemoinformatics: Basic Concepts and Methods. (Wiley-VCH, Weinheim, 2018).

7. Makurvet, F. D. Biologics vs. small molecules: Drug costs and patient access. Med. Drug Discov. 9, 100075 (2021).

8. Lonberg, N. Human antibodies from transgenic animals. Nat. Biotechnol. 23, 1117–1125 (2005).

9. Sharma, P., Joshi, R. V., Pritchard, R., Xu, K. & Eicher, M. A. Therapeutic Antibodies in Medicine. Molecules 28, 6438 (2023).

10. Jin, S. et al. Emerging new therapeutic antibody derivatives for cancer treatment. Signal Transduct. Target. Ther. 7, 39 (2022).

11. Wang, Z. et al. Development of therapeutic antibodies for the treatment of diseases. Mol. Biomed. 3, 35 (2022).

12. Egli, M. & Manoharan, M. Chemistry, structure and function of approved oligonucleotide therapeutics. Nucleic Acids Res. 51, 2529–2573 (2023).

13. Belgrad, J., Fakih, H. H. & Khvorova, A. Nucleic Acid Therapeutics: Successes, Milestones, and Upcoming Innovation. Nucleic Acid Ther. 34, 52–72 (2024).

14. Adachi, T. & Nakamura, Y. Aptamers: A Review of Their Chemical Properties and Modifications for Therapeutic Application. Molecules 24, 4229 (2019).

15. Ellington, A. D. & Szostak, J. W. In vitro selection of RNA molecules that bind specific ligands. Nature 346, 818–822 (1990).

16. Tuerk, C. & Gold, L. Systematic evolution of ligands by exponential enrichment: RNA ligands to bacteriophage T4 DNA polymerase. Science 249, 505–510 (1990).

17. Ng, E. W. M. et al. Pegaptanib, a targeted anti-VEGF aptamer for ocular vascular disease. Nat. Rev. Drug Discov. 5, 123–132 (2006).

18. Danzig, C. J., Khanani, A. M. & Loewenstein, A. C5 inhibitor avacincaptad pegol treatment for geographic atrophy: A comprehensive review. Immunotherapy 1–12 (2024) doi:10.1080/1750743X.2024.2368342.

19. Malicki, S. et al. Identification and characterization of aptameric inhibitors of human neutrophil elastase. J. Biol. Chem. 299, 104889 (2023).

20. Dupont, D. M., Andersen, L. M., Botkjaer, K. A. & Andreasen, P. A. Nucleic acid aptamers against proteases. Curr. Med. Chem. 18, 4139–4151 (2011).

21. Yu, H. et al. Aptameric hirudins as selective and reversible EXosite-ACTive site (EXACT) inhibitors. Nat. Commun. 15, 3977 (2024).

22. Bouhedda, F. et al. A dimerization-based fluorogenic dye-aptamer module for RNA imaging in live cells. Nat. Chem. Biol. 16, 69–76 (2020).

23. Autour, A. et al. Fluorogenic RNA Mango aptamers for imaging small non-coding RNAs in mammalian cells. Nat. Commun. 9, 656 (2018).

24. Hoetzel, J. & Suess, B. Structural Changes in Aptamers are Essential for Synthetic Riboswitch Engineering. J. Mol. Biol. 434, 167631 (2022).

25. Kaiser, C., Schneider, J., Groher, F., Suess, B. & Wachtveitl, J. What defines a synthetic riboswitch? – Conformational dynamics of ciprofloxacin aptamers with similar binding affinities but varying regulatory potentials. Nucleic Acids Res. 49, 3661–3671 (2021).

26. Bouhedda, F., Cubi, R., Baudrey, S. & Ryckelynck, M. μIVC-Seq: A Method for Ultrahigh-Throughput Development and Functional Characterization of Small RNAs. in Small Non-Coding RNAs (ed. Rederstorff, M.) vol. 2300 203–237 (Springer US, New York, NY, 2021).

27. Autour, A., Bouhedda, F., Cubi, R. & Ryckelynck, M. Optimization of fluorogenic RNA-based biosensors using droplet-based microfluidic ultrahigh-throughput screening. Methods 161, 46–53 (2019).

28. Trachman, R. J. et al. Structure and functional reselection of the Mango-III fluorogenic RNA aptamer. Nat. Chem. Biol. 15, 472–479 (2019).

29. Page, M. I. & Badarau, A. The mechanisms of catalysis by metallo beta-lactamases. Bioinorg. Chem. Appl. 2008, 576297 (2008).

30. Hong, D. J. et al. Epidemiology and Characteristics of Metallo-β-Lactamase-Producing *Pseudomonas aeruginosa*. Infect. Chemother. 47, 81 (2015).

31. Ventola, C. L. The antibiotic resistance crisis: part 1: causes and threats. P T Peer-Rev. J. Formul. Manag. 40, 277–283 (2015).

32. Mojica, M. F., Rossi, M.-A., Vila, A. J. & Bonomo, R. A. The urgent need for metallo-β-lactamase inhibitors: an unattended global threat. Lancet Infect. Dis. 22, e28–e34 (2022).

33. Pichon, M. & Hollenstein, M. Controlled enzymatic synthesis of oligonucleotides. Commun. Chem. 7, 138 (2024).

34. Valero, J., et al. A serum-stable RNA aptamer specific for SARS-CoV-2 neutralizes viral entry. Proc. Natl. Acad. Sci. 118, e2112942118 (2021).

35. Zlokarnik, G. et al. Quantitation of Transcription and Clonal Selection of Single Living Cells with β-Lactamase as Reporter. Science 279, 84–88 (1998).

36. Van Berkel, S. S. et al. Assay Platform for Clinically Relevant Metallo-β-lactamases. J. Med. Chem. 56, 6945–6953 (2013).

37. Coleman, T. M. & Huang, F. RNA-Catalyzed Thioester Synthesis. Chem. Biol. 9, 1227–1236 (2002).

38. Stoddard, C. D. et al. Nucleotides Adjacent to the Ligand-Binding Pocket are Linked to Activity Tuning in the Purine Riboswitch. J. Mol. Biol. 425, 1596–1611 (2013).

39. Khan, N. H. et al. A DNA aptamer reveals an allosteric site for inhibition in metallo-β-lactamases. PLOS ONE 14, e0214440 (2019).

40. Wu, D., Gordon, C. K. L., Shin, J. H., Eisenstein, M. & Soh, H. T. Directed Evolution of Aptamer Discovery Technologies. Acc. Chem. Res. 55, 685–695 (2022).

41. Ruckman, J. et al. 2′-Fluoropyrimidine RNA-based Aptamers to the 165-Amino Acid Form of Vascular Endothelial Growth Factor (VEGF165). J. Biol. Chem. 273, 20556–20567 (1998).

42. Hopkins, A. L. & Groom, C. R. The druggable genome. Nat. Rev. Drug Discov. 1, 727–730 (2002).

## References

43. Boussebayle, A. et al. Next-level riboswitch development—implementation of Capture-SELEX facilitates identification of a new synthetic riboswitch. Nucleic Acids Res. 47, 4883– 4895 (2019).

44. Burgstaller, P. & Famulok, M. Isolation of RNA Aptamers for Biological Cofactors by In Vitro Selection. Angew. Chem. Int. Ed. Engl. 33, 1084–1087 (1994).

45. Burmeister, P. E. et al. Direct In Vitro Selection of a 2′-O-Methyl Aptamer to VEGF. Chem. Biol. 12, 25–33 (2005).

46. The pandas development team. pandas-dev/pandas: Pandas. Zenodo 10.5281/ZENODO.10697587 (2024).

47. The Matplotlib Development Team. Matplotlib: Visualization with Python. Zenodo 10.5281/ZENODO.10661079 (2024).

48. Waskom, M. seaborn: statistical data visualization. J. Open Source Softw. 6, 3021 (2021).

49. Bailey, T. L., Johnson, J., Grant, C. E. & Noble, W. S. The MEME Suite. Nucleic Acids Res. 43, W39–W49 (2015).

50. Sievers, F. et al. Fast, scalable generation of high-quality protein multiple sequence alignments using Clustal Omega. Mol. Syst. Biol. 7, 539 (2011).

51. Cock, P. J. A. et al. Biopython: freely available Python tools for computational molecular biology and bioinformatics. Bioinformatics 25, 1422–1423 (2009).

52. Talevich, E., Invergo, B. M., Cock, P. J. & Chapman, B. A. Bio.Phylo: A unified toolkit for processing, analyzing and visualizing phylogenetic trees in Biopython. BMC Bioinformatics 13, 209 (2012).

53. Hagberg, A., Swart, P. J. & Schult, D. A. Exploring network structure, dynamics, and function using NetworkX. (2008).

54. Eaton, D. A. R. Toytree: A minimalist tree visualization and manipulation library for Python. Methods Ecol. Evol. 11, 187–191 (2020).

55. Pedregosa, F. et al. Scikit-learn: Machine Learning in Python. J. Mach. Learn. Res. 12, 2825– 2830 (2011).

56. Crooks, G. E., Hon, G., Chandonia, J.-M. & Brenner, S. E. WebLogo: A Sequence Logo Generator: Figure 1. Genome Res. 14, 1188–1190 (2004).

57. Lee, W., Tonelli, M. & Markley, J. L. NMRFAM-SPARKY: enhanced software for biomolecular NMR spectroscopy. Bioinformatics 31, 1325–1327 (2015).

58. Plateau, P. & Gueron, M. Exchangeable proton NMR without base-line distorsion, using new strong-pulse sequences. J. Am. Chem. Soc. 104, 7310–7311 (1982).

59. Piotto, M., Saudek, V. & Sklenár, V. Gradient-tailored excitation for single-quantum NMR spectroscopy of aqueous solutions. J. Biomol. NMR 2, 661–665 (1992).

60. Sklenář, V., Piotto, M., Leppik, R. & Saudek, V. Gradient-tailored water suppression for 1H-15N HSQC experiments optimized to retain full sensitivity. J Magn Reson 102, 241–245 (2002).

