## Supplementary Material for "Ultrahigh-throughput discovery of modified aptamers as specific and potent enzyme inhibitors"

**Supplementary Table 1| Clade frequency in enriched libraries from the different rounds of selection and screening of the study.**

| Clade | R1 | R2 | R3 | R4 | R5 | R6 | R7 | R7.1 | R7.2 | R7.3 | F7 | F8 | F9 | F10 | F11 | F12 | F12.1 | F12.2 | F12.3 |
|---|---|---|---|---|---|---|---|---|---|---|---|---|---|---|---|---|---|---|---|
| 1 | 0.019 | 0.021 | 0.018 | 0.024 | 0.135 | 0.346 | 0.16 | 0.164 | 0.094 | 0.106 | 0.463 | 0.531 | 0.405 | 0.262 | 0.179 | 0.114 | 0.03 | 0.011 | 0.001 |
| 2 | 0.041 | 0.048 | 0.054 | 0.057 | 0.077 | 0.13 | 0.687 | 0.659 | 0.427 | 0.644 | 0.152 | 0.105 | 0.038 | 0.017 | 0.009 | 0.013 | 0.003 | 0.001 | 0 |
| 3 | 0.02 | 0.021 | 0.019 | 0.027 | 0.309 | 1.281 | 1.414 | 1.506 | 1.437 | 1.479 | 0.705 | 0.29 | 0.07 | 0.026 | 0.016 | 0.02 | 0.01 | 0.006 | 0.001 |
| 4 | 0.175 | 0.212 | 0.246 | 0.293 | 0.9 | 1.272 | 0.305 | 0.298 | 0.139 | 0.18 | 1.129 | 0.634 | 0.238 | 0.101 | 0.046 | 0.044 | 0.017 | 0 | 0 |
| 5 | 0.022 | 0.025 | 0.02 | 0.02 | 0.016 | 0.005 | 0.003 | 0 | 0.001 | 0.002 | 0.005 | 0.003 | 0.002 | 0.002 | 0.002 | 0.001 | 0.109 | 0.383 | 0.514 |
| 6 | 0.018 | 0.019 | 0.019 | 0.022 | 0.018 | 0.006 | 0.004 | 0.003 | 0.002 | 0.001 | 0.005 | 0.003 | 0.003 | 0.002 | 0.002 | 0.001 | 0.218 | 0.728 | 0.809 |
| 7 | 0.012 | 0.017 | 0.015 | 0.018 | 0.097 | 0.399 | 0.685 | 2.802 | 19.249 | 5.087 | 0.396 | 0.264 | 0.099 | 0.041 | 0.014 | 0.107 | 0.012 | 0.011 | 0.003 |
| 8 | 0.067 | 0.082 | 0.064 | 0.086 | 0.182 | 0.506 | 6.232 | 6.151 | 4.644 | 6.892 | 1.781 | 3.13 | 3.571 | 2.766 | 2.295 | 11.976 | 3.847 | 0.838 | 0.239 |
| 9 | 0.462 | 0.674 | 0.542 | 0.709 | 2.103 | 6.627 | 2.449 | 2.659 | 1.612 | 3.501 | 20.733 | 33.126 | 35.24 | 26.23 | 19.932 | 4.422 | 1.324 | 0.2 | 0.038 |
| 10 | 0.008 | 0.007 | 0.007 | 0.005 | 0.004 | 0.001 | 0.003 | 0 | 0.001 | 0 | 0.001 | 0.001 | 0.001 | 0.001 | 0.001 | 0 | 0.258 | 0.919 | 1.173 |
| 11 | 0.166 | 0.187 | 0.171 | 0.183 | 0.176 | 0.141 | 0.1 | 0.095 | 0.05 | 0.083 | 0.27 | 0.698 | 1.971 | 2.896 | 3.547 | 2.611 | 0.721 | 0.151 | 0.055 |
| 12 | 0.018 | 0.024 | 0.021 | 0.026 | 0.037 | 0.097 | 1.592 | 1.55 | 1.375 | 1.522 | 0.117 | 0.096 | 0.055 | 0.027 | 0.023 | 0.068 | 0.034 | 0.015 | 0.001 |
| 13 | 0.02 | 0.025 | 0.023 | 0.028 | 0.21 | 1.261 | 0.692 | 0.729 | 0.421 | 0.755 | 1.029 | 0.632 | 0.262 | 0.095 | 0.064 | 0.03 | 0.013 | 0.006 | 0.001 |
| 14 | 0.042 | 0.045 | 0.042 | 0.046 | 0.37 | 1.939 | 2.311 | 2.274 | 1.768 | 1.651 | 0.782 | 0.219 | 0.042 | 0.021 | 0.016 | 0.021 | 0.005 | 0.01 | 0.013 |
| 15 | 0.259 | 0.319 | 0.237 | 0.298 | 0.292 | 0.293 | 0.054 | 0.049 | 0.033 | 0.05 | 0.624 | 2.901 | 10.289 | 18.182 | 19.124 | 12.332 | 3.818 | 0.754 | 0.191 |
| 16 | 0.034 | 0.043 | 0.038 | 0.043 | 0.208 | 0.603 | 9.142 | 8.583 | 5.679 | 7.561 | 0.543 | 0.262 | 0.08 | 0.026 | 0.011 | 0.1 | 0.051 | 0.019 | 0.004 |
| 17 | 0.065 | 0.084 | 0.067 | 0.106 | 2.154 | 7.935 | 2.629 | 2.817 | 1.495 | 2.412 | 5.093 | 1.839 | 0.435 | 0.141 | 0.08 | 0.026 | 0.01 | 0.008 | 0.001 |
| 18 | 0.028 | 0.041 | 0.029 | 0.051 | 0.766 | 1.748 | 0.764 | 0.684 | 0.341 | 0.613 | 1.762 | 1.083 | 0.31 | 0.1 | 0.041 | 0.03 | 0.009 | 0.004 | 0 |
| 19 | 0.017 | 0.015 | 0.015 | 0.015 | 0.077 | 0.308 | 0.375 | 0.349 | 0.984 | 0.355 | 0.23 | 0.125 | 0.045 | 0.022 | 0.011 | 0.022 | 0.016 | 0.005 | 0.002 |
| 20 | 0.441 | 0.487 | 0.406 | 0.462 | 0.452 | 0.454 | 0.473 | 0.221 | 0.503 | 0.291 | 0.962 | 2.478 | 6.158 | 11.279 | 22.012 | 34.995 | 78.852 | 92.885 | 95.431 |
| 21 | 1.034 | 4.588 | 7.351 | 9.107 | 8.614 | 2.103 | 0.382 | 0.342 | 0.208 | 0.257 | 1.693 | 1.143 | 0.729 | 0.675 | 0.519 | 0.233 | 0.075 | 0.024 | 0.008 |
| 22 | 0.545 | 0.562 | 0.576 | 0.607 | 1.301 | 2.102 | 1.2 | 1.285 | 1.878 | 1.344 | 1.704 | 1.194 | 0.58 | 0.404 | 0.301 | 0.14 | 0.058 | 0.085 | 0.045 |
| 23 | 0.012 | 0.013 | 0.013 | 0.013 | 0.06 | 0.262 | 0.314 | 0.738 | 1.192 | 0.803 | 0.165 | 0.074 | 0.019 | 0.007 | 0.005 | 0.007 | 0.003 | 0.002 | 0 |
| 24 | 0.012 | 0.015 | 0.014 | 0.017 | 0.059 | 0.277 | 0.46 | 0.724 | 3.488 | 1.079 | 0.281 | 0.215 | 0.125 | 0.081 | 0.065 | 0.083 | 0.024 | 0.003 | 0.001 |
| 25 | 0.053 | 0.055 | 0.052 | 0.059 | 0.478 | 2.363 | 3.615 | 3.228 | 2.142 | 3.05 | 1.048 | 0.391 | 0.086 | 0.031 | 0.023 | 0.033 | 0.015 | 0.012 | 0.003 |
| 26 | 0.054 | 0.062 | 0.058 | 0.068 | 0.276 | 1.094 | 1.472 | 1.442 | 1.07 | 1.383 | 1.415 | 1.317 | 0.896 | 0.581 | 0.278 | 0.258 | 0.189 | 0.39 | 0.428 |
| 27 | 0.413 | 0.433 | 0.43 | 0.455 | 0.55 | 0.575 | 0.227 | 0.238 | 0.16 | 0.289 | 0.849 | 1.494 | 1.92 | 1.529 | 0.938 | 0.621 | 0.191 | 0.029 | 0.001 |
| 28 | 0.086 | 0.123 | 0.107 | 0.148 | 1.946 | 9.193 | 17.727 | 16.494 | 11.346 | 14.799 | 6.802 | 3.4 | 0.848 | 0.268 | 0.127 | 0.203 | 0.079 | 0.05 | 0.012 |
| 29 | 0.034 | 0.044 | 0.045 | 0.057 | 0.53 | 1.91 | 1.706 | 1.6 | 0.971 | 1.425 | 1.047 | 0.421 | 0.111 | 0.041 | 0.023 | 0.02 | 0.009 | 0.013 | 0.003 |
| 30 | 0.455 | 0.704 | 0.482 | 0.656 | 0.67 | 0.63 | 0.001 | 0.001 | 0 | 0 | 0.722 | 0.682 | 0.645 | 0.632 | 0.632 | 0 | 0 | 0 | 0 |
| 31 | 0.169 | 0.144 | 0.136 | 0.116 | 0.117 | 0.118 | 0.964 | 0.993 | 0.802 | 0.965 | 0.201 | 0.159 | 0.067 | 0.033 | 0.018 | 0.019 | 0.007 | 0.002 | 0 |
| 32 | 0.085 | 0.12 | 0.092 | 0.137 | 2.139 | 8.545 | 16.332 | 15.319 | 10.661 | 16.155 | 9.466 | 5.16 | 1.513 | 0.558 | 0.209 | 0.277 | 0.072 | 0.036 | 0.012 |
| 33 | 0.094 | 0.095 | 0.091 | 0.094 | 0.705 | 3.441 | 3.314 | 3.243 | 1.909 | 2.654 | 0.682 | 0.145 | 0.043 | 0.03 | 0.027 | 0.027 | 0.012 | 0.01 | 0.001 |
| 34 | 0.624 | 0.657 | 0.585 | 0.595 | 0.507 | 0.312 | 0.044 | 0.042 | 0.033 | 0.034 | 0.33 | 0.306 | 0.262 | 0.258 | 0.252 | 0.008 | 0.007 | 0.019 | 0.026 |
| 35 | 0.71 | 0.588 | 0.522 | 0.439 | 0.326 | 0.133 | 0.024 | 0.024 | 0.015 | 0.011 | 0.069 | 0.053 | 0.044 | 0.044 | 0.039 | 0.002 | 0.002 | 0.001 | 0 |
| 36 | 0.87 | 0.819 | 0.817 | 0.769 | 0.595 | 0.192 | 0.021 | 0.014 | 0.013 | 0.017 | 0.149 | 0.113 | 0.092 | 0.083 | 0.081 | 0.004 | 0.003 | 0.001 | 0 |
| 37 | 1.287 | 1.519 | 1.805 | 1.931 | 1.67 | 0.467 | 0.053 | 0.047 | 0.027 | 0.042 | 0.322 | 0.213 | 0.155 | 0.134 | 0.115 | 0.012 | 0.004 | 0 | 0 |
| 38 | 0.634 | 0.725 | 0.848 | 0.906 | 0.812 | 0.266 | 0.041 | 0.034 | 0.024 | 0.029 | 0.179 | 0.113 | 0.077 | 0.062 | 0.054 | 0.009 | 0.001 | 0 | 0 |
| 39 | 0.009 | 0.013 | 0.014 | 0.016 | 0.027 | 0.049 | 0.085 | 0.303 | 1.332 | 0.396 | 0.031 | 0.012 | 0.003 | 0.002 | 0.002 | 0.008 | 0 | 0.001 | 0 |
| 40 | 0.058 | 0.062 | 0.059 | 0.063 | 0.282 | 1.116 | 1.287 | 1.21 | 0.729 | 1.11 | 0.674 | 0.304 | 0.068 | 0.027 | 0.016 | 0.018 | 0.005 | 0.004 | 0.001 |
| 41 | 0.016 | 0.018 | 0.021 | 0.025 | 0.069 | 0.21 | 0.231 | 0.449 | 2.683 | 0.916 | 0.202 | 0.148 | 0.058 | 0.028 | 0.014 | 0.032 | 0.005 | 0.001 | 0 |
| 42 | 0.079 | 0.112 | 0.108 | 0.137 | 0.934 | 4.597 | 5.315 | 5.158 | 3.513 | 4.143 | 4.169 | 2.626 | 0.799 | 0.267 | 0.135 | 0.11 | 0.048 | 0.015 | 0.003 |
| 43 | 0.033 | 0.036 | 0.046 | 0.05 | 0.055 | 0.074 | 1.055 | 0.98 | 0.653 | 0.746 | 0.084 | 0.059 | 0.025 | 0.014 | 0.011 | 0.021 | 0.008 | 0.003 | 0 |
| 44 | 0.021 | 0.025 | 0.021 | 0.028 | 0.237 | 1.162 | 1.451 | 2.262 | 7.051 | 2.717 | 0.617 | 0.286 | 0.081 | 0.031 | 0.015 | 0.049 | 0.009 | 0.006 | 0.015 |
| 45 | 0.104 | 0.121 | 0.104 | 0.121 | 0.381 | 0.808 | 0.565 | 0.503 | 0.393 | 0.57 | 0.956 | 0.951 | 0.597 | 0.36 | 0.197 | 0.179 | 0.069 | 0.015 | 0.012 |
| 46 | 0.262 | 0.26 | 0.246 | 0.243 | 0.223 | 0.164 | 0.072 | 0.063 | 0.033 | 0.059 | 0.195 | 0.35 | 0.53 | 0.603 | 0.493 | 0.269 | 0.09 | 0.037 | 0.092 |
| 47 | 0.191 | 0.282 | 0.233 | 0.317 | 0.896 | 3.649 | 3.861 | 3.647 | 1.74 | 4.559 | 6.66 | 12.391 | 15.07 | 15.081 | 11.94 | 10.693 | 3.444 | 0.604 | 0.156 |

**Supplementary Table 2 | Main sequences used throughout the study.**

| Name | Sequence |
|---|---|
| Starting library | 5’-GGATCCGACCGTGGTGCCNNNNNNNNNNNNNNNNNNNNNNNNNNNNNNNNNNNNNNNNGCAGTGAAGGCTGAGCTCC-3’ |
| Fwd | 5’-AATTC**TAATACGACTCACTATA**GGAGCTCAGCCTTCACTGC-3’ |
| Ampli-Forward | 5’-AATTC**TAATACGACTCACTATA**GGGAGACTCAGCCTTCAC-3’ |
| Ampli-Reverse | 5’-GGATCCGACCGTGGTGCC-3’ |
| Add-barcode primer | 5’-TCGTCGGCAGCGTCAGATGTGTATAAGAGACAGAANNNNNNNNNNNNNNNNNNNNAATTC**TAATACGACTCACTATA**GGGAGACTCAGCCTTCAC-3’ |
| Ampli-Barcode | 5’-TCGTCGGCAGCGTCAGATGTGTATAAGAGA-3’ |
| NGS-Reverse | 5‘-GTCTCGTGGGCTCGGAGATGTGTATAAGAGACAGGGATCCGACCGTGGTGCC-3‘ |
| S1 | 5’-GGGAGACTCAGCCTTCACTGCTAGAGATTCGGTGGGACACCCGGTGAAAGGCTGACCACTAGGCACCACGGTCGGATCC-3’ |
| S1.1** | 5’-GGGATAATTCAGCCTTCACTGCTAGAGATTCGGTGGGACACCCGGTGAAAGGCTGAATTATCC-3’ |
| S1.2** | 5’-GGGATAATTCAGCCTTCACUUCGGTGAAAGGCTGAATTATCC-3’ |
| S1.3** | 5’-GGGATAATTCAGCCTTCACTGCTAGAGATTCCCCGGGACGGGCGGTGAAAGGCTGAATTATCC-3’ |
| S1.4** | 5’-GGGATAATTCAGCCTTCACTGCTAGAGATTCCCCGUUCGCGGGCGGTGAAAGGCTGAATTATCC-3’ |
| S1.5** | 5’-GGGATAATCCTGCTAGAGATTCGGTGGGACACCCGGATTATCCC-3‘ |
| S2 | 5’-GGGAGACTCAGCCTTCACTGCTAGAGATTCGGTGGGACACCCTGTGAAAGGCTGACCACTAGGCACCACGGTCGGATCC-3’ |
| S25** | 5’-GGGAGACGGGTGTCCCACCGAATCTCTAGCATCTCCC**TATAGTGAGTCGTATTA**GAATTGTGA-3’ |
| S30** | 5’-GGGAGAAGGGTGTCCCACCGAATCTCTAGCATCTCCC**TATAGTGAGTCGTATTA**GAATT-3’ |
| 34** | 5’-GGGAGATGGGTGTCCCACCGAATCTCTAGCATCTCCC**TATAGTGAGTCGTATTA**GAATT-3’ |
| 35** | 5’-GGGAGACGGGTGTCCCACCGAATCTCTAGCGTCTCCC**TATAGTGAGTCGTATTA**GAATT-3’ |
| 36** | 5’-GGGAGAAGGGTGTCCCACCGAATCTCTAGCCTCTCCC**TATAGTGAGTCGTATTA**GAATT-3’ |
| 39** | 5’-GGGAGAAGGGTGTCCCACCGAATCTCTAGCTTCTCCC**TATAGTGAGTCGTATTA**GAATT-3’ |
| 40** | 5’-GGGAGAGGGGTGTCCCACCGAATCTCTAGCCTCTCCC**TATAGTGAGTCGTATTA**GAATT-3’ |
| 41** | 5’-GGGAGACGGGTGTCCCACCGAATCTCTAGCTTCTCCC**TATAGTGAGTCGTATTA**GAATT-3’ |
| 42** | 5’-GGGAGATGGGTGTCCCACCGAATCTCTAGCCTCTCCC**TATAGTGAGTCGTATTA**GAATT-3’ |
| 32** | 5’-GGGAGACGGGTGTCCCACCGAATCTCTTGCATCTCCC**TATAGTGAGTCGTATTA**GAATT-3’ |
| 38** | 5’-GGGAGAAGGGTGTCCCACCGAATCTCTTGC ATCTCCC**TATAGTGAGTCGTATTA**GAATT-3’ |

* T7 RNA polymerase promoter or its reverse complement is bolded. ** Template synthesized in reverse complement orientation.

**Supplementary Table 3 | Inhibition and interaction parameters of SPM1 inhibitory aptamers.**

| | | **Inhibition capacity** | | | **Affinity** | | | |
|---|---|---|---|---|---|---|---|---|
| | | Imipenem | CCF2-FA | CENTA | BLI | | | EMSA |
| | | $k_{iapp}$ (nM) | $k_{iapp}$ (nM) | $k_{iapp}$ (nM) | $K_D$ (nM) | $k_{on}$ (x $10^5$ M-1s-1) | $k_{off}$ (x $10^{-3}$ s-1) | $K_D$ (pM) |
| S1 | RNA | 162 ± 22 | 73 ± 40 | 516 ± 34 | 28 ± 2 | 1.55 ± 0.15 | 4.37 ± 0.13 | n.d. |
| | 2'-FY | 38 ± 10 | <23 | n.d. | 24 ± 2 | 0.68 ± 0.06 | 1.63 ± 0.29 | n.d. |
| S25 | RNA | 289 ± 33 | 64 ± 10 | n.d. | 53 ± 7 | 1.88 ± 0.09 | 9.98 ± 1.10 | 298 ± 78 |
| | 2'-FY | 58 ± 8 | <15 | 279 ± 60 | 54 ± 12 | 1.17 ± 0.11 | 6.18 ± 0.81 | 131 ± 24 |
| S30 | RNA | 193 ± 23 | n.d. | n.d. | 20 ± 3 | 1.66 ± 0.14 | 3.36 ± 0.65 | 77 ± 12 |
| | 2'-FY | 48 ± 7 | n.d. | 112 ± 37 | 38 ± 1 | 0.86 ± 0.07 | 3.28 ± 0.34 | 165 ± 3 |

**a**

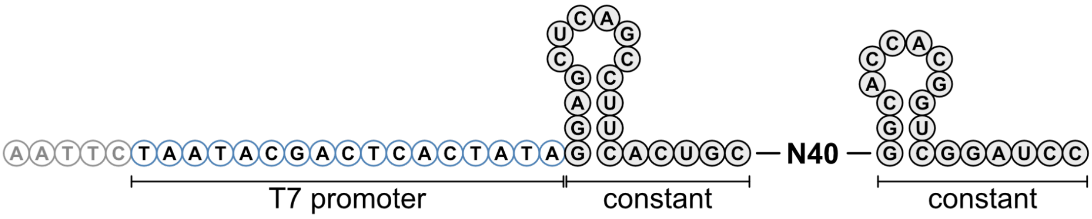


**b**

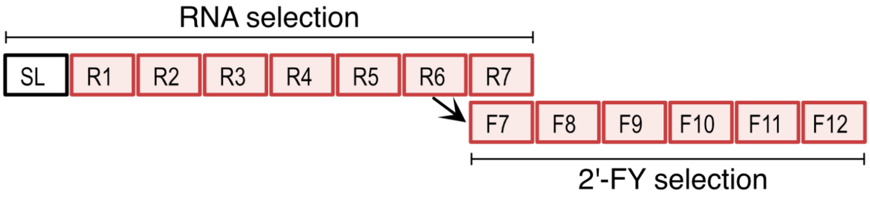


**c**

| Round | Chemistry | % eluted from target | % eluted from beads | Amount SPM-1 (nmol) | Amount RNA (nmol) |
|---|---|---|---|---|---|
| R1 | RNA | 0.18 | 0.11 | 0.5 | 2 |
| R2 | RNA | 0.15 | 0.07 | 0.5 | 2 |
| R3 | RNA | 0.12 | 0.09 | 0.5 | 2 |
| R4 | RNA | 0.15 | 0.08 | 0.5 | 2 |
| R5 | RNA | 0.38 | 0.3 | 0.5 | 2 |
| R6 | RNA | 3.27 | 0.39 | 0.5 | 2 |
| R7 | RNA | 9.91 | 0.14 | 0.5 | 2 |
| F7 | 2'-FY | 0.23 | 0.1 | 0.5 | 2 |
| F8 | 2'-FY | 0.3 | 0.17 | 0.5 | 2 |
| F9 | 2'-FY | 0.36 | 0.11 | 0.5 | 2 |
| F10 | 2'-FY | 1.24 | 0.2 | 0.5 | 2 |
| F11 | 2'-FY | 1.29 | 0.21 | 0.5 | 2 |
| F12 | 2'-FY | 9.33 | 2.89 | 0.5 | 2 |

**Supplementary Fig. 1 | *In vitro* selection (SELEX) experiments. a**, Design of the starting library used in the study. T7 RNA polymerase promoter (blue circles) was placed upstream the construct to drive its transcription. This promoter sequence was present in the starting library and reintroduced after every round of selection during RT-PCR. The pool of RNA or 2'-FY oligonucleotides (black circles) was then obtained by *in vitro* transcription. Note that the 40-nucleotide long randomized region (N40) was flanked by constant region folding as small stem-loops that prestructure the pool. **b**, *In vitro* selection workflow. The starting library (SL) was first expressed in RNA and subjected to 7 rounds of SELEX (R1-R7). The enriched pool obtained from round R6 was then expressed in 2'-FY chemistry and subjected to 6 additional rounds of SELEX (F7 to F12). **c**, Summary table of the SELEX process. Third and fourth column: Amount of RNA recovered after selection from SPM-1 coupled beads or empty beads, respectively.

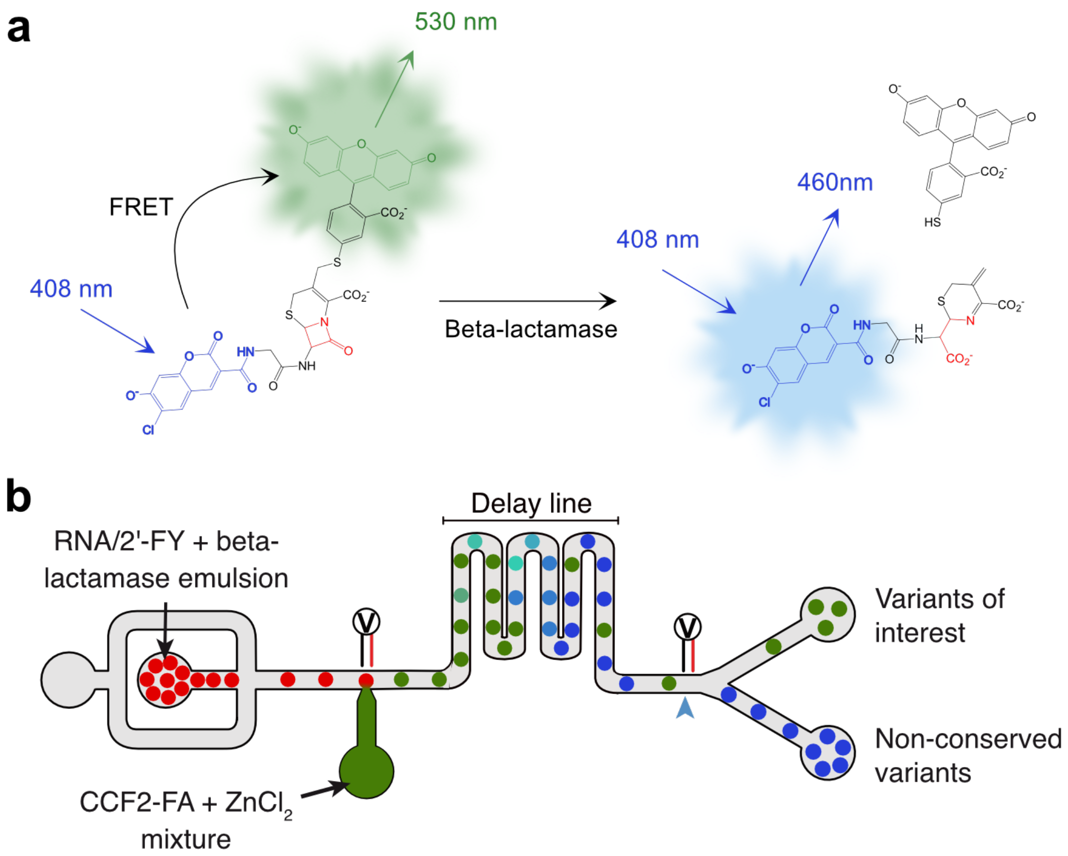


**Supplementary Fig. 2 | Microfluidic-based beta-lactamase activity assay. a**, Fluorescent CCF2-FA beta-lactamase activity assay. CCF2-FA is a beta-lactamase substrate analogue exploiting fluorescence resonance energy transfer (FRET) consisting of a blue-emitting B7-hydroxycoumarin-linked to a green emitting fluorescein via a cephalosporin core[1]. Upon excitation at 408 nm, the coumarin moiety of the uncleaved substrate excites the fluorescein, leading to green fluorescence emission. Upon cephalosporin cleavage by a beta-lactamase, fluorescein is released from coumarin, and blue fluorescence is emitted upon excitation at 408 nm. **b.** Integrated microfluidic chip device for high-throughput enzyme activity assay in droplets. Droplets containing the transcripts (RNA or 2'-FY) and the enzyme are reinjected into the device and spaced by an oil stream. An aliquot of fluorescent substrate (CCF2-FA) and zinc chloride ($ZnCl_2$) is then delivered to each droplet by picoinjection. Next, the droplets enter a delay line at controlled temperature to enable the enzyme to transform its substrate. Finally, the droplets reach the end of the device where their blue and green fluorescences are individually measured (blue arrow). The blue/green fluorescence ratio is then computed to evaluate enzyme activity and sort the droplets accordingly by applying an electric field.

1. Zlokarnik, G. et al. Quantitation of Transcription and Clonal Selection of Single Living Cells with β-Lactamase as Reporter. Science **279**, 84–88 (1998).

**a**

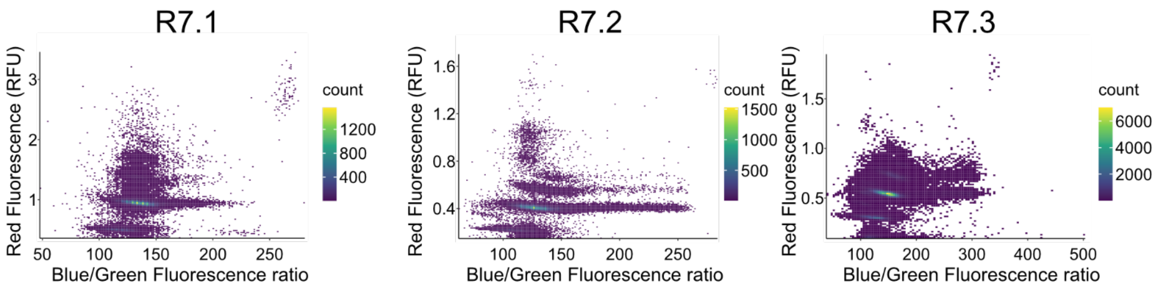


| | R7.1 | R7.2 | R7.3 |
|---|---|---|---|
| Number of analyzed droplets | 1 923 214 | 2 823 261 | 2 777 989 |
| Number of sorted droplets | 14 993 | 35 867 | 72 550 |
| % of sorted droplets | 0.77 | 1.27 | 2.6 |

**b**

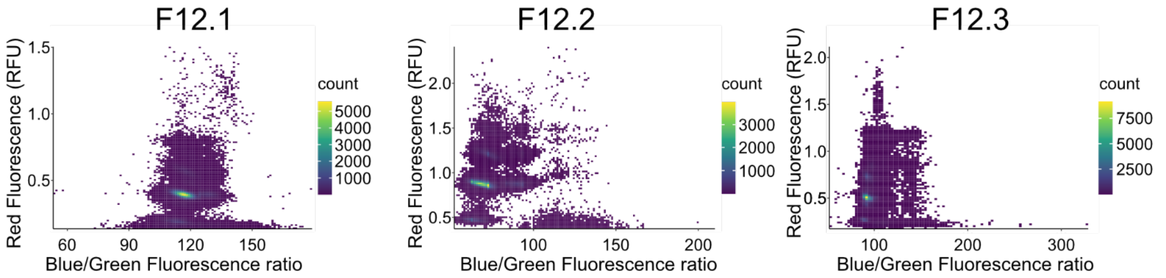


| | F12.1 | F12.2 | F12.3 |
|---|---|---|---|
| Number of analyzed droplets | 2 178 614 | 1 539 711 | 1 234 711 |
| Number of sorted droplets | 112 332 | 46 583 | 14 439 |
| % of sorted droplets | 5 | 3 | 1.2 |

**c**

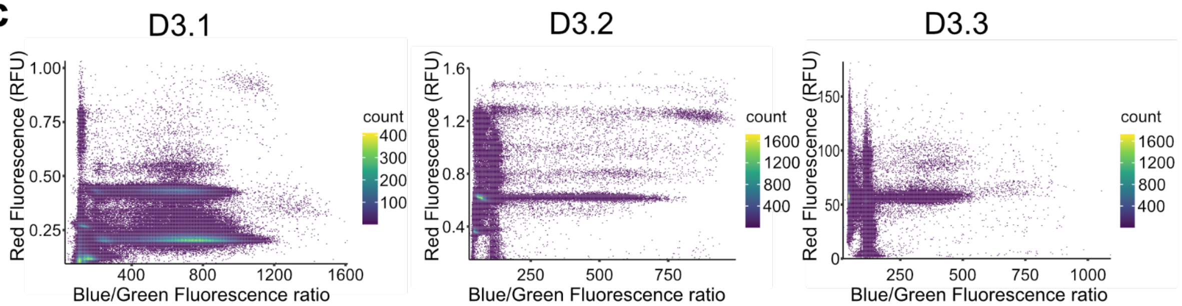


| | D 3.1 | D 3.2 | D 3.3 |
|---|---|---|---|
| Number of analyzed droplets | 2282992 | 785983 | 2286108 |
| Number of sorted droplets | 35989 | 2177 | 35989 |
| % of sorted droplets | 1.6 | 0.8 | 1.6 |

**Supplementary Fig. 3 | Fluorescence profiles and statistics of emulsions analyzed during the rounds of screening.** The red (Cy5, droplet tracer), blue and green (CCF2-FA) fluorescence of each droplet was measured. Red fluorescence allows to discriminate unfused *in vitro* transcription droplets (low Cy5 fluorescence) from those fused with one PCR droplet (intermediate Cy5 fluorescence) or more than one PCR droplet (high Cy5 fluorescence). Blue/green fluorescence ratio was computed to determine the CCF2-FA conversion: the higher the ratio that higher the beta-lactamase activity. **a**, RNA screening, **b**, 2’-FY screening, **c**, Doped pool screening.

**a**

| Clade # | Motif | RNA | 2'-FY |
|---|---|---|---|
| 1 | ACBDDWWKHMYGWCHRSHGDSCHCGYKGTSWCHAVAHYB | - | + |
| 2 | YGYKRTCRACRAWKMKSGMWRMWMMGMRRMSCTAWYMSK | + | - |
| 3 | GRKGCCCSGWRCKSTWRWKAMARCGKAWYMARSCKSWWY | + | + |
| 4 | CCGAAACTTCCTACTACTAAGATATACCCGTGCAACCTG | + | + |
| 5 | TTAGGTCATGCTCGAATGACCGTGCTGCATTAAGCGGAA | - | + |
| 6 | TCAGGCTATTCTACTGCTCTACCGTGATGAACGAGCAGA | - | + |
| 7 | GATATACTGGCGGTAGCCGACAATGAAATGTAACGCTAA | + | - |
| 8 | TGCTATCAACGAATCGGGCAGCACCGCGGAGCTATTCGT | + | + |
| 9 | TGMTATCAAMGAATCRSDYADMMCCKMSGAGCKRYKCKK | + | + |
| 10 | GGGAAAGGACATCCGGGACGGATTTGAGAGAAAGTTTCT | - | + |
| 11 | AATGGGGAATCCCCGCACTATAGAGATA | - | + |
| 12 | CAAGGTGCCTTATGTCAAACTAGAGCGATACGACCTCTG | + | - |
| 13 | CWWGSYGMCWKRYKKYWAAMTAGMGMSATACRRYMTSYG | + | + |
| 14 | CTTGCCGACAGGCTGTTAAATAGCGACCTACAGTAGGCG | + | + |
| 15 | GATCCTAGACGGGGAGGTATACGGTGGTTGCATTTAAAT | - | + |
| 16 | CCAGCTCGAGCGCGCTAAAACGCTCAAACCACGCTACGA | + | + |
| 17 | CMAGSTMGAGMSCGCYWARAMGCTSMAAMSACKYYWSRA | + | + |
| 18 | CAAGGTAGAGACCGCCTAGAAGCTGMAAAGACTTCTGAA | + | + |
| 19 | AGADATWCKGBSGGWMRCCSAWWRWRARRYSKAMCRCTA | + | - |
| 20 | TAGAGATTCGGTGGGACACCCGGTGAAAGGCTGACCACT | + | + |
| 21 | AAHAAWAAYAAYAAHAAMMM | + | + |
| 22 | AHACTRHAGAGAHAACDCGWCYDWSVAG | + | + |
| 23 | SAYMRTGWAAWSTRRMGMKAMRRCASGAKKKTGKWSRSA | + | - |
| 24 | ACATCGTGAAAACTGGAGAGACAACACGATTGTGTAGGC | + | - |
| 25 | GCGTTGCGGGGACTGGATTATCAAATCACACCGAGGAAG | + | + |
| 26 | CTGTTTTACTGTCAGCAGAGCTCGTTGTCACAAGAACCA | + | + |
| 27 | CAATACGGGAATAGTGCTTCCGAGCGATCCACCAT | + | + |
| 28 | CGTGGTCGACAATGATCGATAATAAGAAACCCTAACACG | + | + |
| 29 | GACGTGTGGTATCATCAGAATATACCCAAACCCTAACAC | + | + |
| 30 | ACCCATGGTGGATCGCTCGGAAGCACTATTCCCGTATTG | + | + |
| 31 | HMDGCTCKAGCVCGSKWRMHCVCYSAWASSWYG | + | - |
| 32 | ATGCTCTAGCACGGGTGCCCACCGATAGGTTGTAAAGAC | + | + |
| 33 | TATCCAACTATCCACGCCCCGCCGGCAGGCTGTTAAGTG | + | + |
| 34 | RRWSGCMGTAACTTCTAGCGCAAGT | + | + |
| 35 | SBCMCBCCMMMCHCMDHNCRCACNCCCC | + | + |
| 36 | YDMACCMSCSCMCNAWMACCVCGCC | + | + |
| 37 | ACAHHAYMHHAYMMHYMCMYYNCCMMCM | + | + |
| 38 | ACTAYAMWYAMGCVCSCCAC | + | + |
| 39 | ACTCGACAATGAAATGTAACGCTTAGATTCAGTCAGTAA | + | - |
| 40 | AGAGACAAGTCCCTGTCACGTAATAMAGGACCAAAGGGC | + | + |
| 41 | AGAGACAACTCTGTGATTTTCTGCGAAATCACAAGCGCA | + | - |
| 42 | AGWTAAMAAAAAACTACATAAACCCTACCACGGGCGCTG | + | + |
| 43 | AASWKAASARAMAACWRSWYWAASSCKRCSASSGRMSCK | + | - |
| 44 | AACTGAAGAGACAACAGGTCTAAGGCGGCGAGCGAACCG | + | + |
| 45 | TCAGAACACGATTCAAATGACGCCCGCGGTTCAAATCA | + | + |
| 46 | AGAGGGTGAACGAAATACACTTTGCCCGCGGTAGCCAGT | - | + |
| 47 | ATAAGATTAACGAATCACWTATGCCCSSGGKKMGSC | + | + |

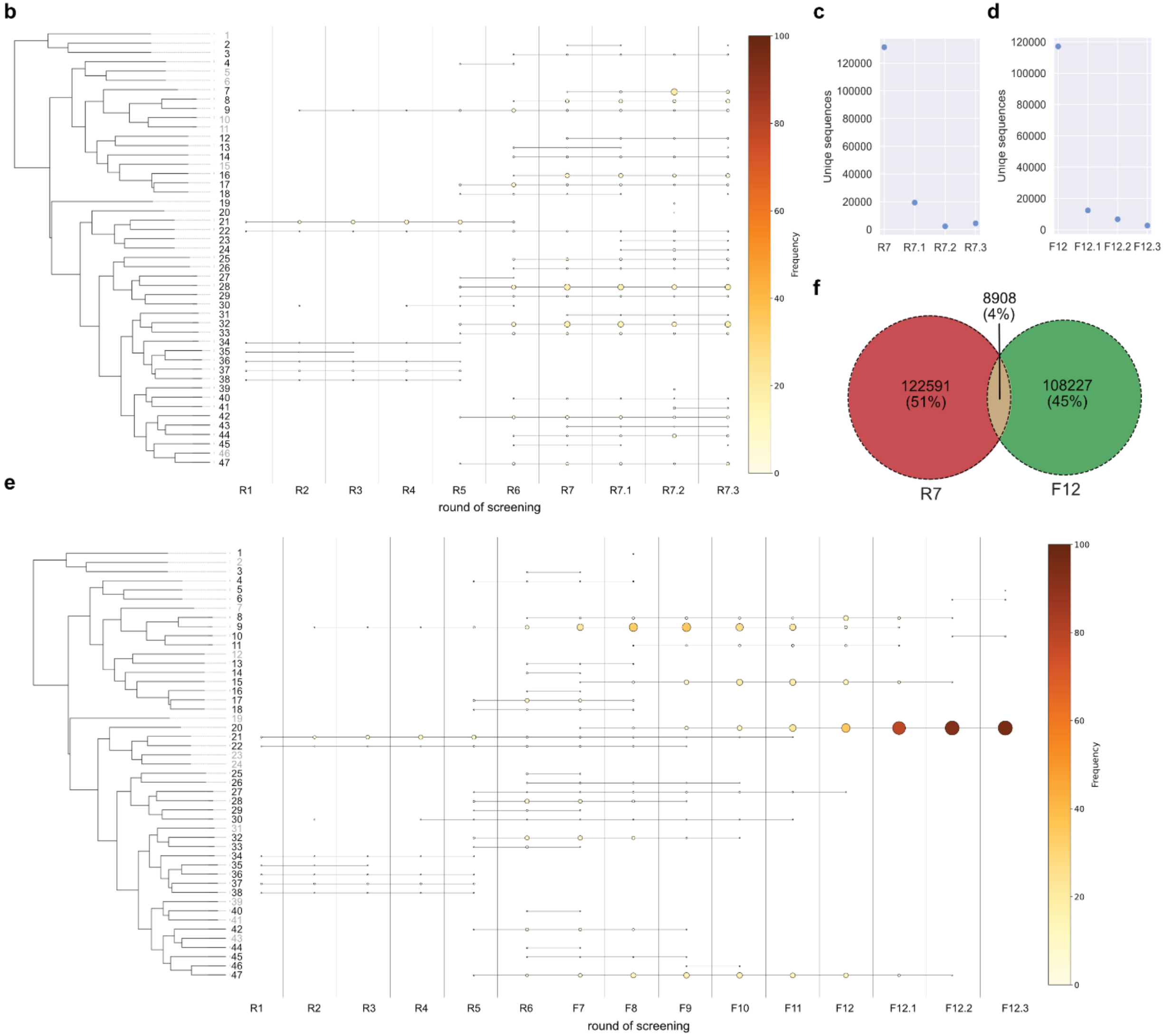


**Supplementary Fig. 4 | Bioinformatic analysis of library content. a**, Representative consensus sequences of the 47 identified enriched clades. **b**, and **e**, Phylogenetic distribution of the clades throughout selection rounds R1 to R7.3 and R1 to F12.3. Clades were organized according to their phylogenetic similarity and their appearance frequency is encoded by the size and the color point. Clades absent for the selection are depicted in a light gray. **c**, and **d**, Number of unique sequences found in libraries R7 and after subsequent rounds of screening, and in F12 and after subsequent rounds of screening. **f**, Venn diagram showing the number of unique sequences found in rounds R7 and F12. The diagram shows the number of unique sequences and their frequency.

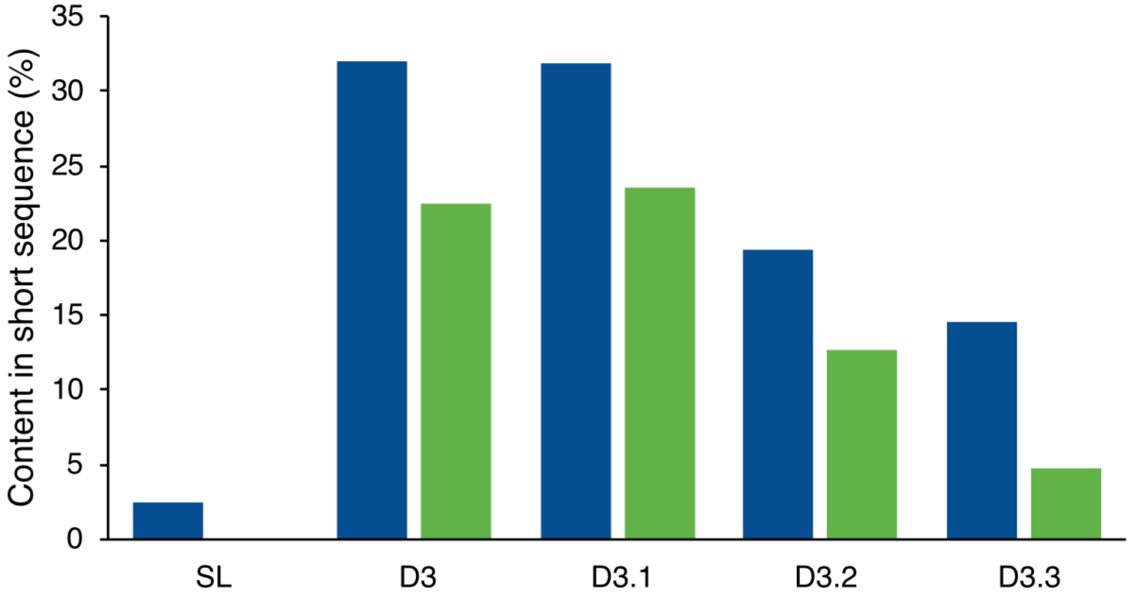


**Supplementary Fig. 5 | Microfluidic-promoted depletion of unwanted small sequences.** The size distribution of the sequences contained in the starting library (SL), those enriched after the last round of doped-SELEX (D3) or after each round of functional screening (D3.1-D3.3) was analyzed using microcapillary electrophoresis (Agilent 2100 Bioanalyzer, blue bars) or determined from sequencing data (green bars). In both cases, the percentage of sequences of the expected size (i.e., 154 nucleotides for the bioanalyzer or 37 nucleotides for the bioinformatics) was calculated.

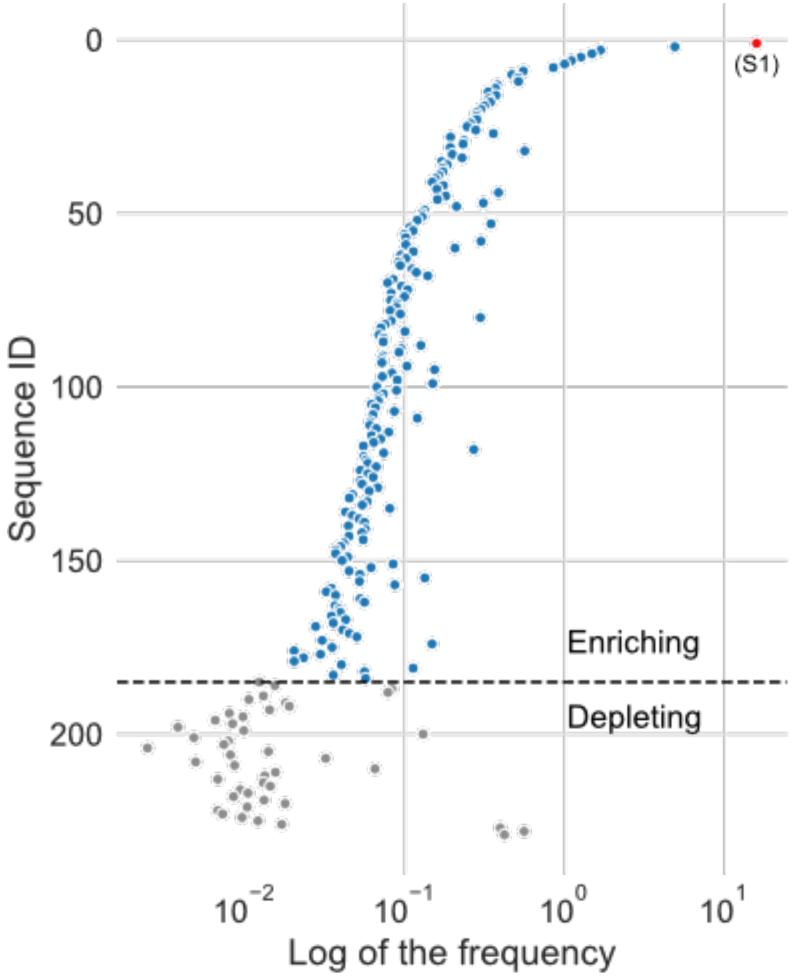


**Supplementary Fig. 6 | Frequency of clade 20 sequences in round F12.3.** Sequences are ordered according to their enrichment throughout rounds F12.1 to F12.3. Sequences displaying an enrichment are symbolized in blue whereas those depleting are symbolized in gray. The dominant variant S1 is highlighted in red.

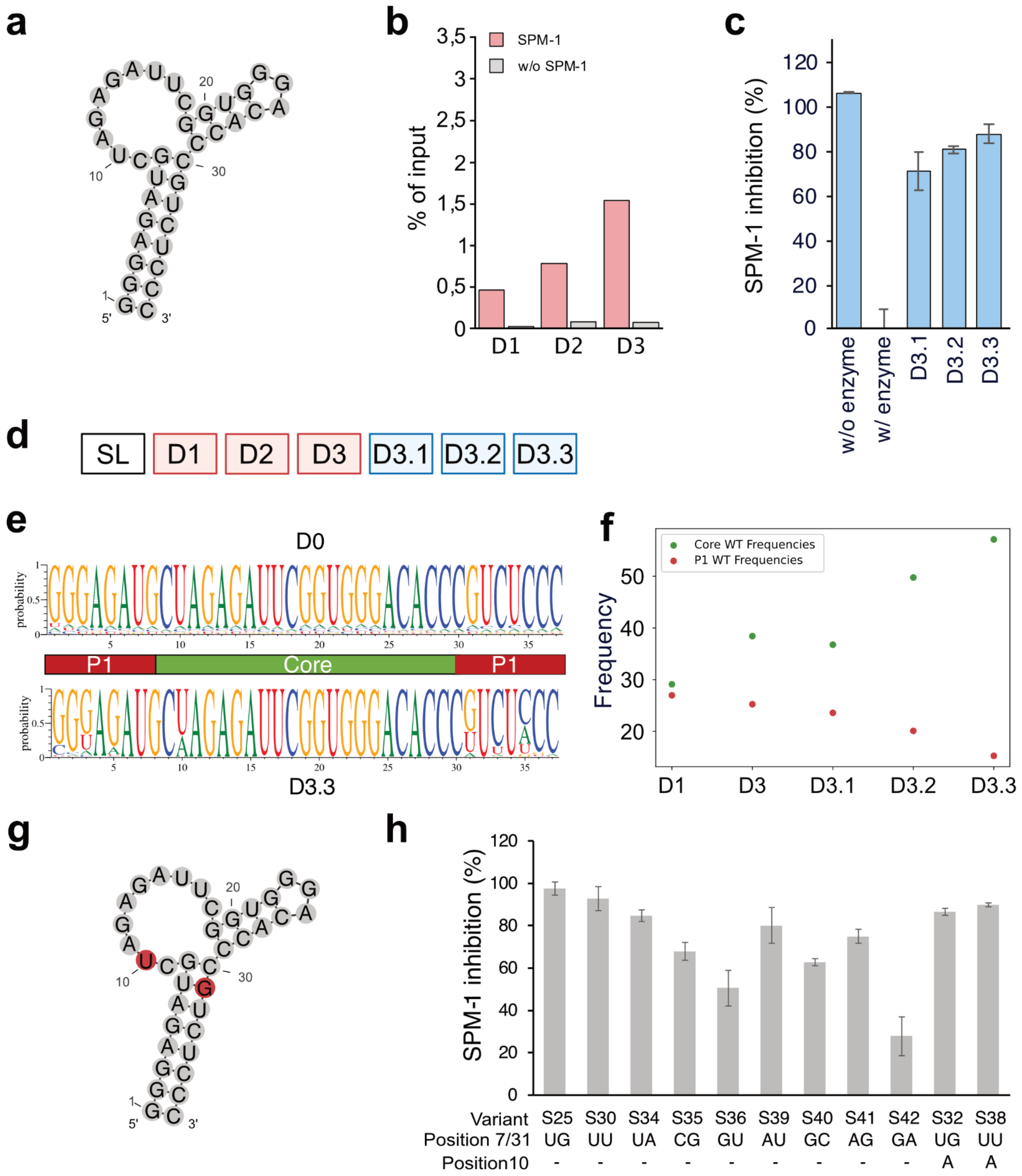


**Supplementary Fig. 7 | Selection, screening and characterization of S25 doped pool. a**, Secondary structure model of the starting library based on S25. Each position was randomized to 9% (3% of each nucleotide). **b**, Elution profile of the oligonucleotide bound fraction during SELEX experiments. The fraction of oligonucleotide eluted from SPM-1-coupled beads (red bars) or empty beads (negative control, gray bars) is given as the percentage of the total input RNA used in each round. **c**, Enrichment profile of the libraries in SPM-1 inhibitory aptamers. Libraries recovered after each round of screening (D3.1-D3.3) were expressed in 2'-FY (light blue) and their capacity to inhibit SPM-1 activity was assessed using CCF2-FA assay. **d**, Doped selection workflow. SELEX steps (D1-3, red boxes) are distinguished from screening steps (D3.1-3.3, blue boxes). SL: starting library. **e**, Base composition of the starting (SL) and the final enriched (D3.3) libraries. P1 stem and core aptamer regions are delineated. **f**, Occurrence frequency of wild-type P1 stem (green dots) and core aptamer (red dots) sequences throughout selection and screening steps. **g**, Secondary structure model of S25 aptamer. The nucleotides showing sequence tolerance on **e** are labeled in red. **h**, Functional validation of mutants identified or designed upon doped selection. Mutations of position 31, its predicted complementary position 7 and position 10 were prepared in 2'-FY and functionally tested using imipenem assay. Values are the mean of 3 independent experiments and the error bars correspond to the standard deviation. Error bars were cut upon axis interception for an illustrative purpose.

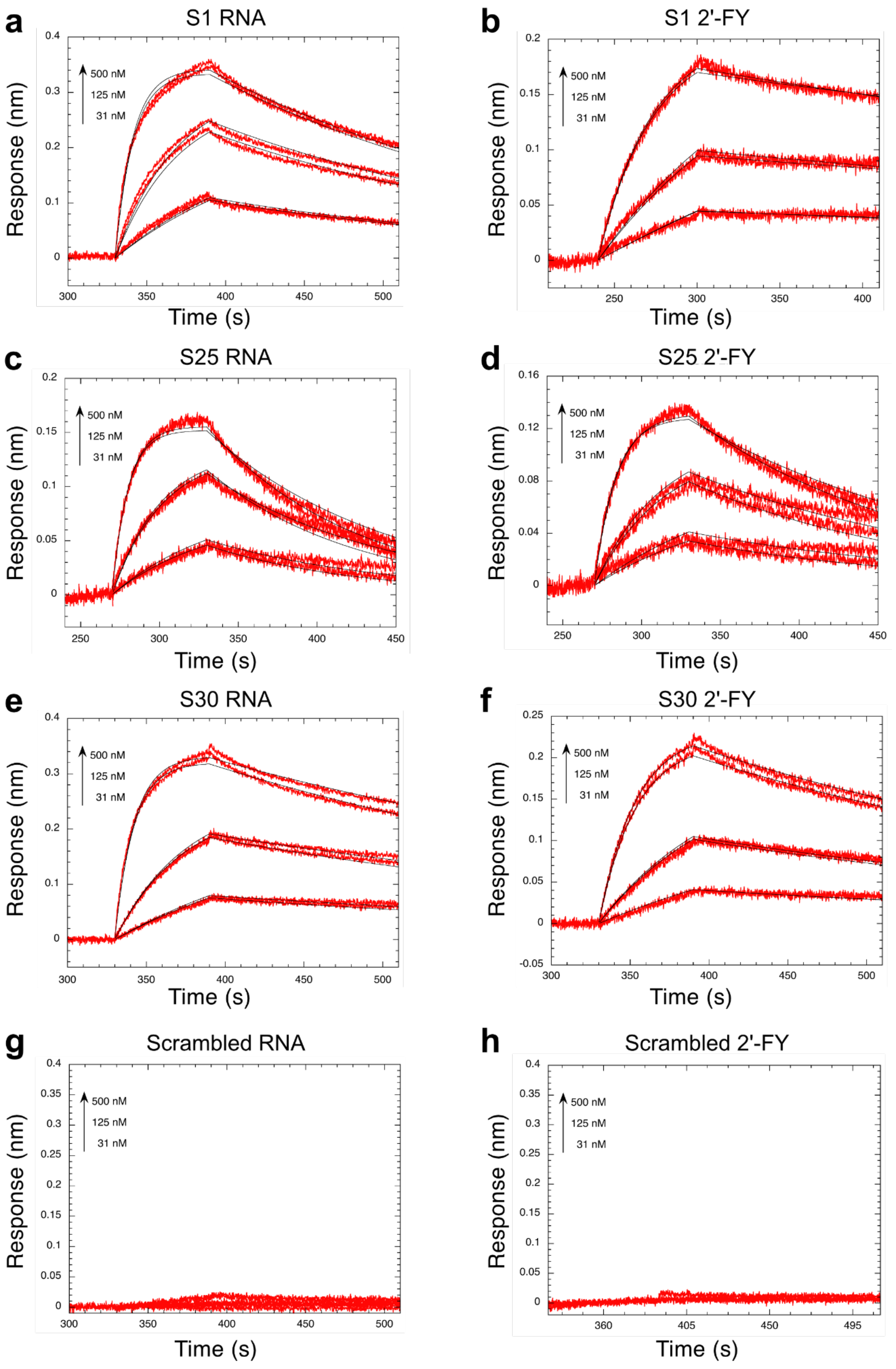


**Supplementary Fig. 8 | Binding kinetics of oligonucleotides to SPM-1 monitored by BLI.** The protein was captured on Ni-NTA BLI sensors and incubated with increasing concentration of oligonucleotide as indicated by the arrows. The red and black curves correspond to the experimental data and the fits to a 1:1 model of interaction, respectively.

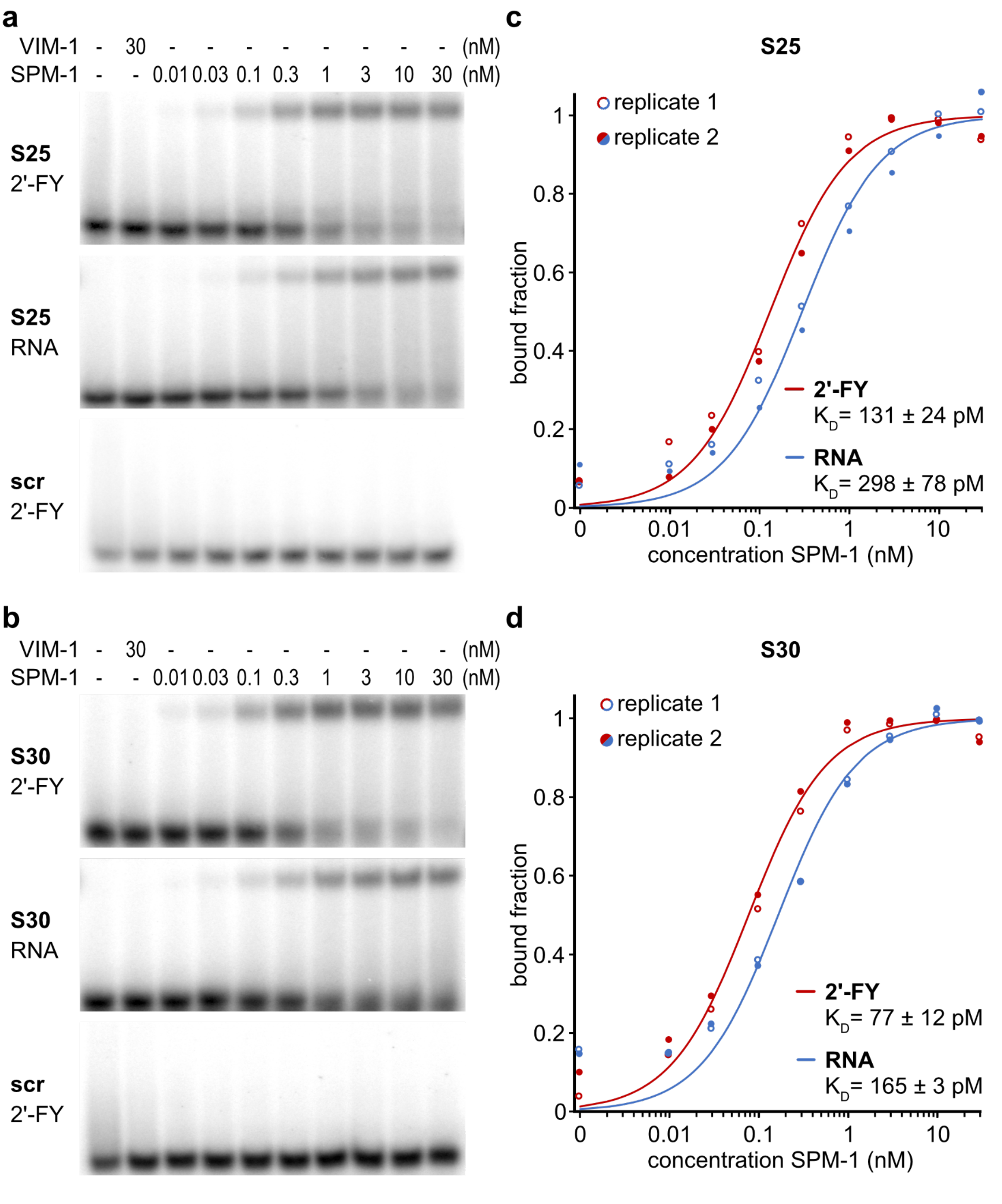


**Supplementary Fig. 9 | Determination of oligonucleotides/SPM-1 binding affinities using EMSA. a**, Electrophoretic mobility shift assay gels using S25 transcribed in 2'-FY or RNA, or a scrambled control (scr). VIM-1 was used as a control protein for unspecific binding. **b**, Electrophoretic mobility shift assay gels of S30 transcribed in 2'-FY or RNA, or a scrambled control (scr). VIM-1 was used as a control protein for unspecific binding. **c**, Curve fitting of the binding of S25 to SPM-1 in 2'-FY and RNA. **d**, Curve fit of the binding of S30 to SPM-1 in 2'-FY and RNA. Dissociation constant ($K_D$) was determined by fitting data to a Hill equation. Values are the mean of two independent replicates.

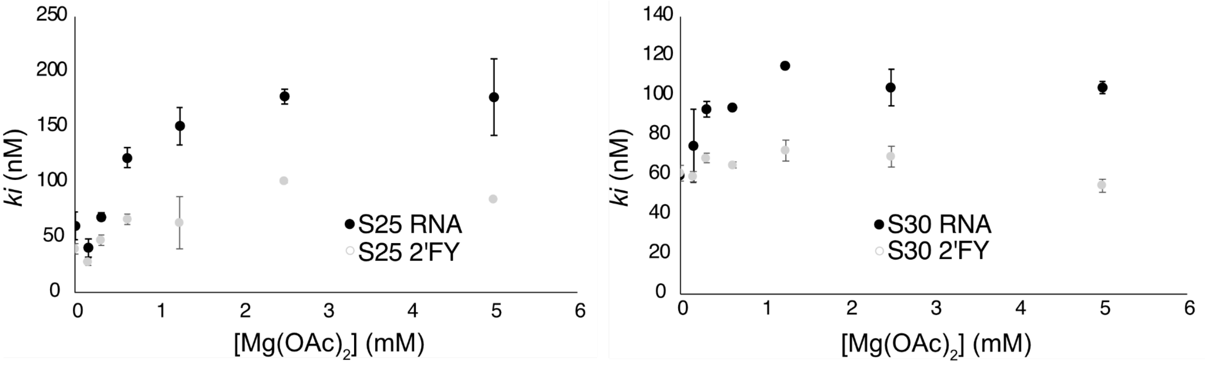


**Supplementary Fig. 10 | Influence of magnesium acetate concentration on aptamer-mediated SPM-1 inhibition.** Inhibition constant ($k_i$) of SPM-1 activity by aptamers in RNA or 2'-FY chemistry was determined over different concentrations of magnesium acetate ($MgOAc_2$).

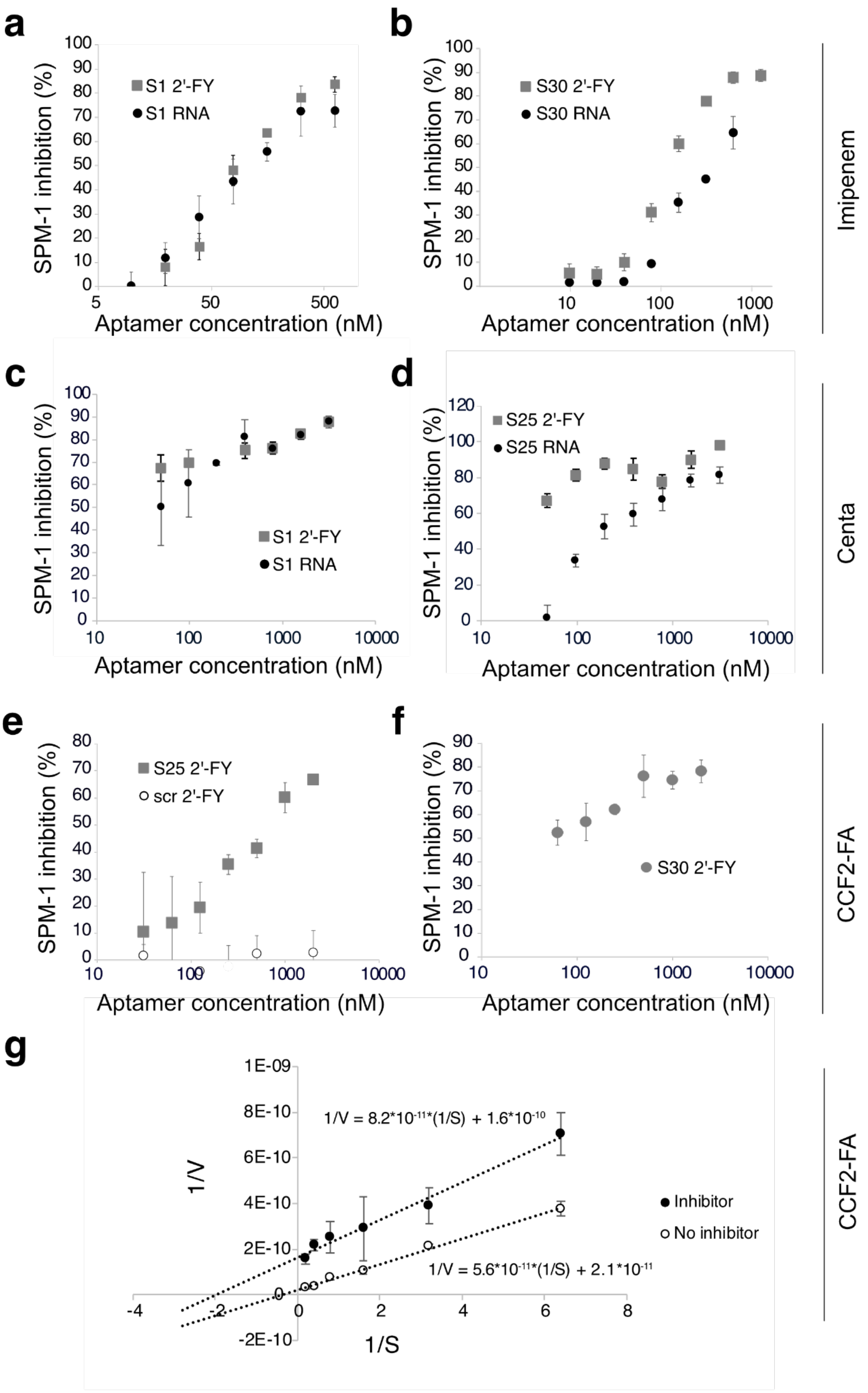


**Supplementary Fig. 11 | SPM-1 inhibition kinetics. a**, Inhibition of SPM-1 activity by S1 and **b**, S30 aptamers evaluated with imipenem assay. **c**, Inhibition of SPM-1 activity by S1 and **d**, S25 aptamers evaluated with CENTA assay. **e**, Inhibition of SPM-1 activity by S25 and **f**, S30 aptamers evaluated with CCF2-FA assay. In most cases, aptamers were evaluated in RNA and 2'-FY chemistries by incubating a range of aptamer concentrations with SPM-1 and monitoring enzyme activity. **g**, Kinetic characterization of aptamer-mediated inhibition of SPI-1 activity. Initial reaction rates were determined at several concentrations of CCF2-FA in the presence of 7 nM nM of SPM-1 and in presence or absence of 50 nM S1 2'-FY aptamer.

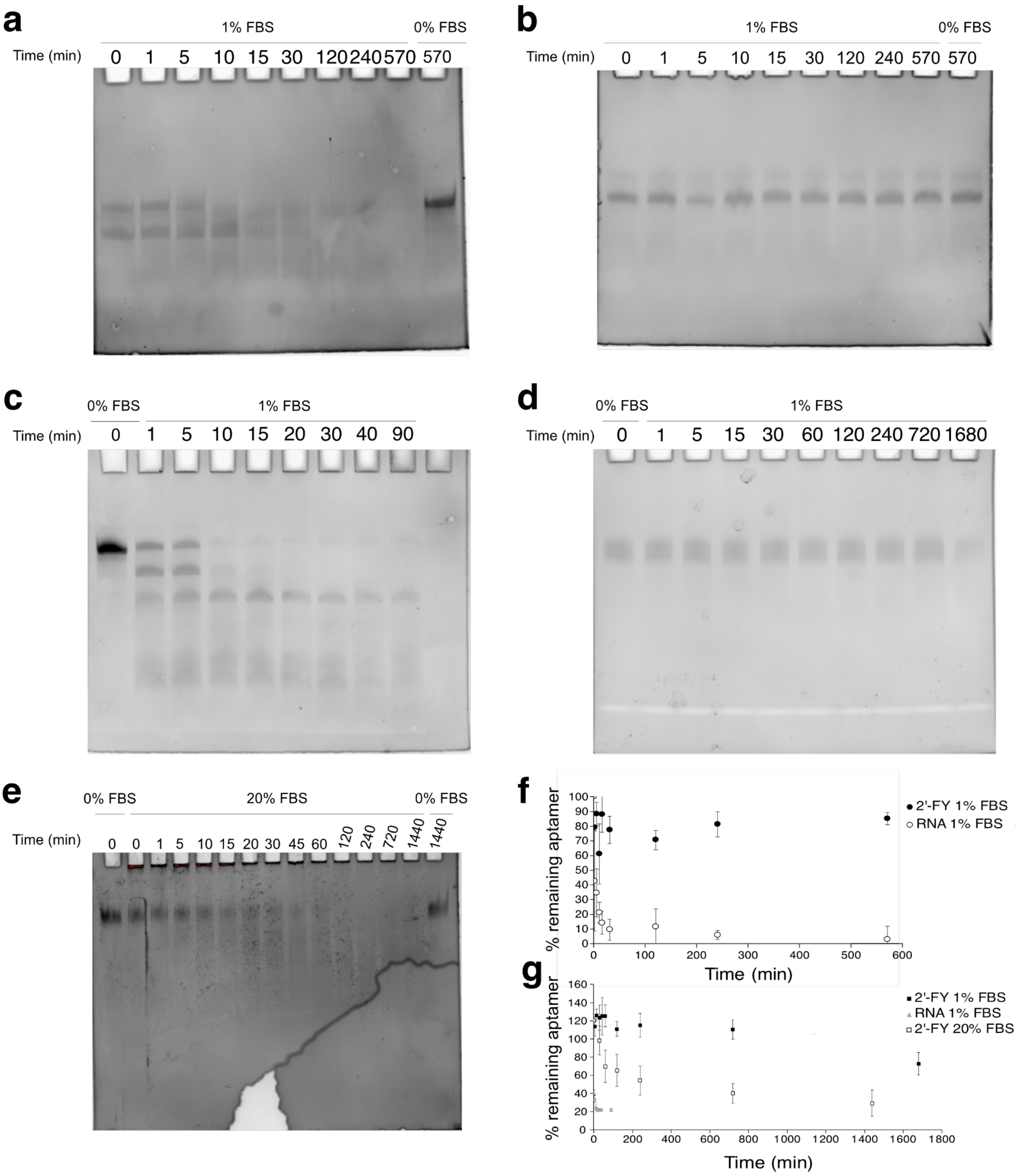


**Supplementary Fig. 12 | Nuclease resistance of aptamers. a**, S25 RNA aptamer degradation kinetics in 1% FBS. **b**, S25 2'-FY aptamer degradation kinetics in 1% FBS. **c**, S30 RNA aptamer degradation kinetics in 1% FBS. **d**, S30 2'-FY aptamer degradation kinetics in 1% FBS. **e**, S30 2'-FY aptamer degradation kinetics in 20% FBS. **f**, Quantification of S25 aptamers stability in 1% FBS. The bands of gels from panels **a** and **b** were quantified by densitometry. **g**, Quantification of S30 aptamers stability in 1% FBS. The bands of gels from panels **c** and **d** were quantified by densitometry.
